# Using the pan-genomic framework for the discovery of genomic islands in the haloarchaeon *Halorubrum ezzemoulense*

**DOI:** 10.1101/2023.10.13.561781

**Authors:** Yutian Feng, Danielle Arsenault, Artemis S. Louyakis, Neta Altman-Price, Uri Gophna, R. Thane Papke, Johann Peter Gogarten

## Abstract

In this study, we use pan-genomics to characterize the organized variability from the widely dispersed halophilic archaeal species *Halorubrum ezzemoulense*. We include a multi-regional sampling of newly sequenced, high-quality draft genomes. Using the pan-genome graph of the species, we discover 50 genomic islands which represent rare accessory genetic capabilities available to members of the species. 19 of these islands are likely the remnant of mobile genetic elements and are enriched for genomic dark matter. 10 islands encode for niche adapting solute transporters, with a cosmopolitan but limited distribution throughout the strains. We also observe rearrangements which have led to the insertion/recombination/replacement of mutually exclusive genomic islands in equivalent genome positions (“homeocassettes”). These conflicting islands encode for similar functions, but homologs from islands located between the same core genes exhibit extreme divergence on the amino acid level. Homeocasettes provide variations for a homologous function, which may confer a greater range of adaptability to the species group. We observe some islands that appear geographically restricted; however, we also observe the coexistence of genomes, in a single geographic region, with and without certain genomic islands, demonstrating the retention and spread of rare genes in the pan-genome.

**Importance:** Understanding the evolution of genome content is a key puzzle in evolutionary biology. Despite its importance, this area hasn’t received thorough investigation. This is especially true of Archaeal organisms, which constitute a large fraction of Earth’s diversity, but are often referred to as the “forgotten” or “third” domain of life. This study dives into those questions by finding rare genes amongst a group of closely related Archaeal species, and describes how their transfer, utilization and persistence may contribute to the speciation and specialization of the group.

## Introduction

Genomic islands (GIs) are loci in a genome that were acquired through horizontal gene transfer. Genes within GIs frequently differ from those in the rest of the genome (1, 2) in coding density, local nucleotide composition and codon usage (2). Acquisition through HGT or recombination is a fundamental part of the GI definition. Compositional differences from the genome’s average, the presence of mobility elements and their proximity to tRNA genes are frequently used as surrogate methods, albeit often ineffective (Ragan, 2001; Cortez et al., 2005), to identify GIs in addition to their limited distribution among a group of related organisms (5, 6). The term “genomic island” is an umbrella term that encompasses different types of islands, including defense, pathogenicity/virulence, and niche adaptation. The typing of a genomic island is determined by the genes encoded on it and the phenotype/capabilities it confers onto the host post-acquisition. Defense islands encode for defense systems such as restriction modification systems, which target foreign DNA, and are active players in the genetic arms race which underlies the evolution of prokaryotic organisms which partake in horizontal gene transfer (7). Pathogenicity islands can fundamentally alter the effect of a microorganism on the disease state of its eukaryotic host (8). Niche adapting genomic islands encode for capabilities which may improve the fitness or survivability of the organism, which harbors it, in response to specific environmental stresses. Genomic islands exhibit diverse origins and functions; based on their persistence, ubiquity, transferability, and encoded functions, they clearly have played decisive roles in the evolution of prokaryotic organisms.

There is no doubt as to the importance of horizontal gene transfer’s (HGT or HT: horizontal transfer) impact on evolution (9, 10). The successful transfer of a functional gene between two extant organisms may instantaneously provide the recipient with novel traits bypassing the drawn-out adaptation of a gene already present in the genome through mutations. This utility extends to genomic islands, as they encompass a set of interacting genes which may encode for a related function (i.e., HGT “en-masse”). The transferability of genomic islands is reflected by their limited distribution in related genomes (members of the same genus/species will not have the same islands), as well as the enrichment of transposases, conjugation related genes and integrases in or around the immediate gene neighborhood of genomic islands (5, 11, 12).

The haloarchaea are one of the best known and well sampled groups of Archaea. Their obligate halophily (often in environments with salt concentration above 2.0M NaCl) (13, 14), selective phototrophy (15), and metabolic capabilities (16, 17) are well documented. In addition to halophily, the genome content of haloarchaea reflects environmental pressures like limited nutrient availability, prolonged exposure to UV radiation, and the genetic arms race with selfish genetic elements and viruses. Haloarchaea are well-known for their ability to exchange DNA within and between species and even across domains (18–20). Haloarchaeal species frequently undergo homologous recombination, facilitating the spread of alleles and genes within local populations (20–24). This characteristic paves the way for the acquisition (from outside the species), and transfer of loci between closely related genomes.

To investigate the gene content and diversity of genomic islands in haloarchaea, we focused on a single species, *Halorubrum ezzemoulense* (*Hez).* This species was first described in 2006 based on a single strain cultivated from an Algerian sabkha (25). Later, with more cultivated strains, *Hez* was evaluated using a polyphasic approach (21) and the species was expanded and eventually amended to include *Halorubrum chaoviator* strains (26). Standard experimental DNA-DNA hybridization (DDH) values of *Hez* were correlated to their 16S rRNA and multi-locus gene sequence divergences (21). In each case, the 70% DDH cut-off corresponded to >99 % nucleotide sequence identity for all the genes analyzed in their sample. We assembled a dataset consisting of 47 isolates of *Halorubrum ezzemoulense*, the DNA of 37 of which were newly sequenced and assembled into draft genomes for this study (see data availability section for the genome files and Table S1 for genome information). Every genome of our current dataset is a member of the same species, having at least 98.5% pairwise average nucleotide identity, though most of the strains share above 99% identity (Table S2). These strains were isolated from various geographic locations, including but not limited to: Santa Pola Saltern (Spain), Aran-Bidgol (Iran), and the Dead Sea (Israel). We note that this multiple isolate approach does not reflect the true diversity or composition of *Hez* in their respective geographic locations; but only represents a limited and possible biased sample of the diversity present in these places. Limiting the dataset to members of the same species informs on what kind of rare/flexible gene content is maintained and diversified in that group.

We report the distribution and content of 50 GIs with a limited distribution within the *Hez* species, discovered using the pan-genome graph approach described in Panaroo (27). 10 of these islands contain the components of solute transporters, demonstrating differential metabolic capabilities within members of the same species. We also find that different GIs often are found inserted between the same core genes and, furthermore, encoding for related functions in a mutually exclusive fashion. We dubbed these matching islands “homeocassettes”. Although inserted between the same core genes and encoding in part homologous genes, these homeocassettes exhibit extreme divergence and are the result of duplication and reassortment events.

## Results and Discussion

### *Hez* pan-genome reveals flexible gene content

Using the Panaroo pipeline, we used sequence identity and local synteny to cluster the predicted genes throughout all 47 *Hez* genomes into pan-gene families. When available, pan-gene families were named according to their annotations’ consistency throughout all the genomes, otherwise pan-gene families were named by arbitrary ascending group numbers. The entire genetic repertoire or “pan-genome” of *Hez* consists of 8,401 pan-genes. The core genome, defined here as those genes found in at least 44 of 47 (~94%) of all *Hez* draft genomes, includes 2,621 pan-gene families (Fig S1). The shell genome, here defined as non-core genes identified in at least 2 genomes, is the largest fraction of the pan-genome (4,072 pan-genes). This may be a consequence of different genetic specializations/mosaicism present in the species and/or reflect predation by mobile genetic elements. Genes which are found in only one genome are considered in this study to be a part of the cloud division of the pan-genome. Pan-genome accumulation and core-genome rarefaction curves, as well as a description of each pan-genome division can be found in Fig S1. Detailed information of the 47 *Hez* genomes including assembly statistics, number of unique pan-genes, and secondary replicon information is included in Table S1. These pan-genome inferences reveal a flexible genome content and produced a gene-connectivity graph used to discover and elucidate the local context of 50 pan-genomic islands.

### Isolates Overview

Pairwise comparisons of total average nucleotide identity (tANI) which incorporate genome alignment fraction (genome distances were determined by −ln(pairwise ANI + alignment fraction)), following methods by Gosselin et al., 2022 (Gosselin et al., 2022), confirm that all 47 strains are of the same species (Table S2). The phylogeny based on corrected tANI data (Fig 1) reveals regionally delimited clans (highlighted clans from Spain and Israel in Fig 1), and well-supported groups formed by isolates from multiple regions. Isolates from Spain and Israel not placed within the regionally delimited groups (Fig 1, highlighted clades) cluster with isolates from other regions. The tANI phylogeny reflects both, sequence variation in the core genome and the flexible gene content across all genomes. Another phylogeny (Fig S2), based only on core genome differences, was inferred using a nucleotide concatenate of all 2,621 core genes (where ~2.2M of 2.3M sites were constant). Both phylogenies recover a similar topology including both regionally delimited groups, but strains not associated with a group were placed differently due to shared accessory genes. For example, strain UG96 shares several genomic islands with other Israeli isolates (Island 1, 4, 23, etc.; Fig 2), and is placed with the Israeli isolates in the tANI phylogeny but is placed outside this group with respect to the core genes. This observation is consistent with the notion that flexible genome content, such as the genes encoded on genomic islands, impact placement of groups in the tANI phylogeny (Fig 1).

**Figure 1.**
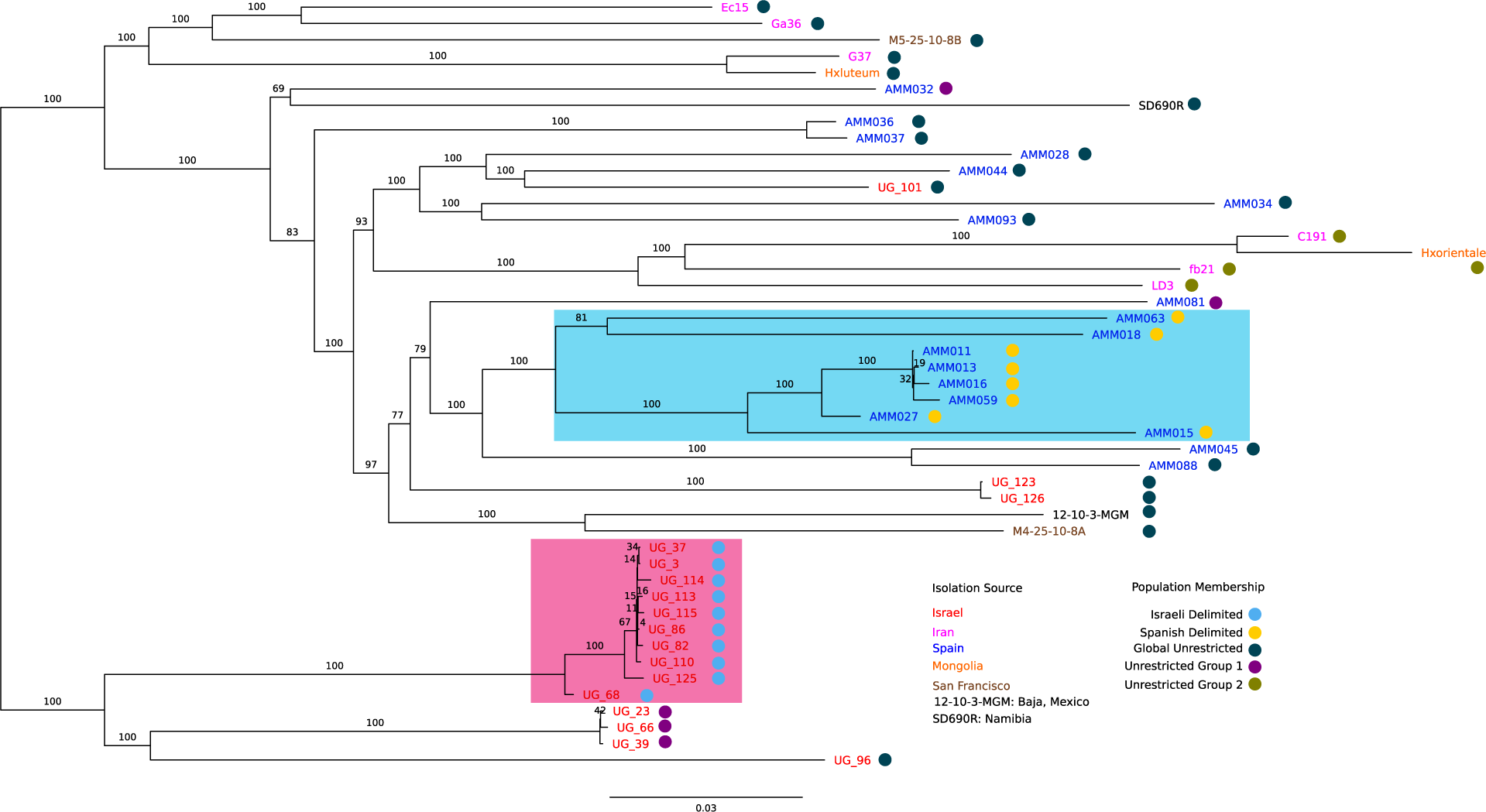
Overview of Hez strain phylogeny and nested population structure. The phylogeny was calculated using pairwise ANI and genome alignment fraction and bootstrapped following [28]. A distance matrix showing all the pairwise relationships can be found in Table S2. Branch supports are denoted by the numbers above the branches. The coloring of the tip labels indicates isolation region. Each strain has been assigned membership into nested populations using rhierBAPs (based on SNPs shared in their selectively neutral core genes, see methods), which is designated by the colored circle at each tip. The highlighted clades indicate that the population membership (reflecting core genome) and phylogenetic grouping (reflecting whole genome) both agree that these are regionally delimited (in the sense that the nested population structure forms a monophyletic group on the phylogeny) clans in Spain (cyan) and Israel (pink).

**Figure 2.**
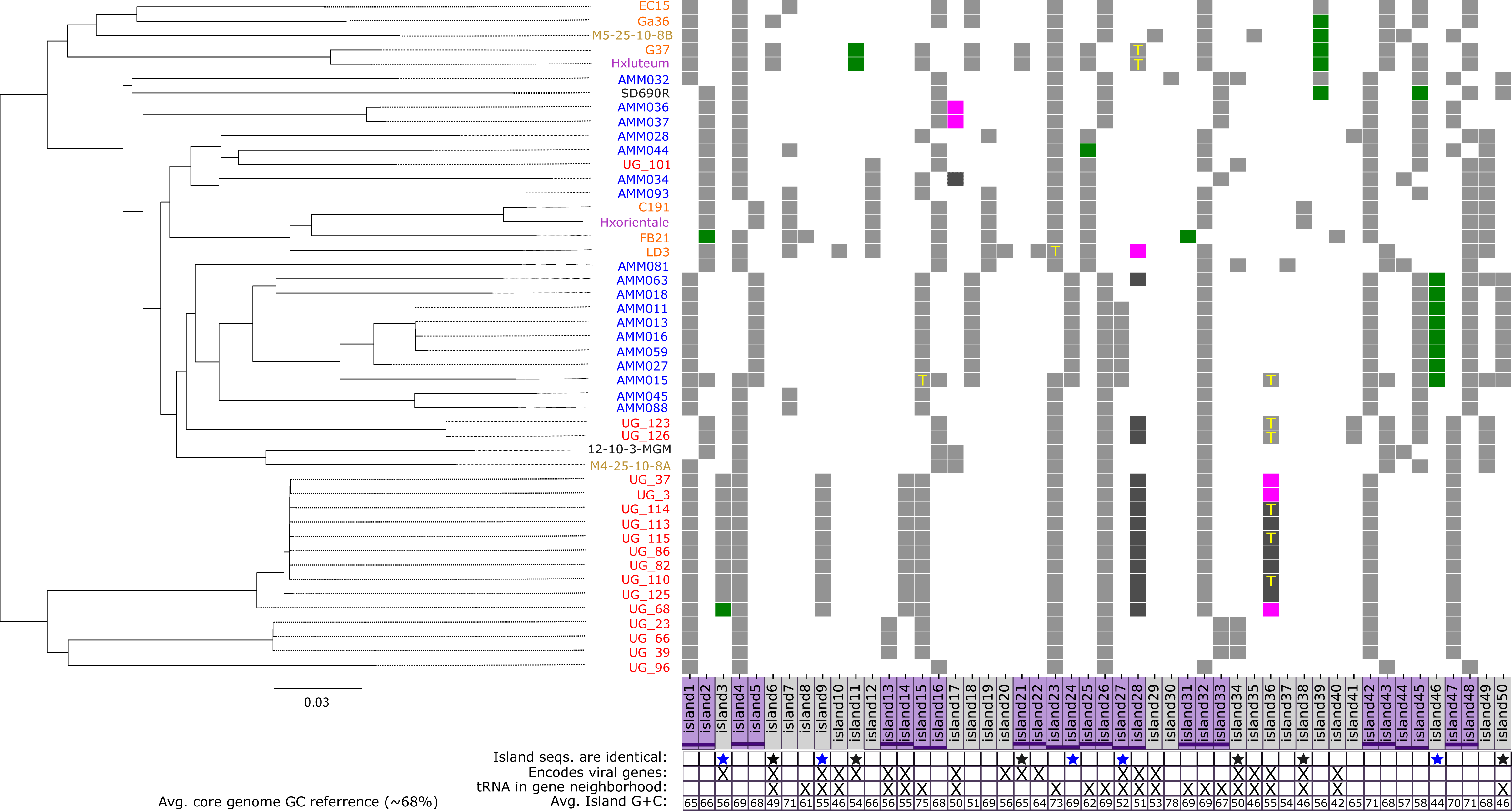
Distribution of genomic islands in *Halorubrum ezzemoulense*. The presence/absence map of discovered pan-genomic islands (x-axis) mapped onto the tANI phylogeny. The y-axis are the genomes in which the islands are present, the tip labels are colored based on the region of isolation (red: Dead Sea, Israel; blue: Santa Pola, Spain; brown: San Francisco, USA; green: Iran; purple, Mongolia; SD690R: Namibia; and 12-10-3-MGM: Baja California, Mexico). The purple boxes indicate conflicting genomic islands, islands invading the same core genes have a purple horizontal dash spanning adjacent purple boxes, at the same height. Below the island name plates is a table which conveys various contexts of each corresponding island. A blue star beneath the island name indicates that the island sequences are identical in the genomes that have it (*i.e.*, Island 27’s sequence is identical across the Spanish strains which have it). Similarly, a black star indicates that the sequences of that island are identical, and the island is found in multiple geographic regions (and coexists in those regions with isolates without the island). Green boxes on the heat-map indicate that a transposase (IS element) was found in the genomic neighborhood (±5 genes) of the island, in that genome. Exact family designations and names of transposases can be found in Table S7. Magenta boxes indicate that a flanking transposase was found but the ORFs on the genomic island have been shuffled or that the island is only a partial representation of the full genomic island found in other genomes. Similarly, dark gray boxes show the partial genomic island without the presence of a flanking transposase.

In addition to phylogenies, we performed hierarchical clustering of isolates using rHierBAPs (Tonkin-Hill et al., 2018) (which is based on SNPs found in neutral core genes, where neutrality was determined using Tajima’s D values <|0.2|). This inference revealed a nested population structure within *Hez* (Fig 1; population membership shown with circle indicators). The regionally delimited clans in Spain and Israel observed in the tANI phylogeny correspond to nested population structures when analyzing the SNPs of neutral core genes (Fig 1, highlighted clans). This may reflect potential barriers to dispersal and recombination in these regions. These regionally delimited groups (Fig 1: blue circles [Israel] and gold circles [Spain]) may represent lineages diverging within the overall Hez species. The inferred populations, excluding those confined to a single clade (purple, dark blue, and olive circles), indicate information exchange or migration at a global scale. The largest population, represented by the dark blue circle (Global), encompasses SNP contributions from strains in every isolation location and spans almost every group in the phylogeny. This may reflect either an ancestral lineage of *Hez* or a cosmopolitan strain with little barriers to recombination.

### Islands in the Hez pan-genome

The Panaroo pipeline produces a pan-genome graph (27), that represents gene families and their connectivity in each genome. The pan-genome graph is analogous to a De Bruijn assembly graph, where the nodes are gene families instead of contigs, and the edges describe the gene adjacency information for each gene family. Node by node traversal of this pan-genome graph, considering the genome representation at each node (i.e., a core gene has a representation = 47 and a unique gene = 1), returns the pan-genomic islands that have been integrated in between core genes (see methods “Querying the Pan-genomic Graph”). By employing this pan-genome graph-based approach, larger scale gene transfers can be reliably detected even when the local gene neighborhood is confounded with gene acquisition/loss, structural rearrangements, and poor quality of the draft genome assembly. Reliability in this case stems from the requisite of core genes flanking the island, as the local gene neighborhood and representation of the core genes must be recovered from at least 44 of 47 *Hez* genomes. This is analogous to the approach which identifies genomic islands as low coverage regions when metagenomic reads are mapped onto an isolate’s genome (30, 31). The pan-genome graph resembles the reference genome, while individual genomes act as metagenomic reads. The sharp drop from high coverage spanning core genes to pan-genomic islands with low coverage illustrates the flexibility of gene content. We only considered pan-genomic islands which encode at least two genes (although islands of size one were included when they were found to invade the same core genes as another island; see below “homeocassettes”).

Analysis of the pan-genome graph reveals 50 distinct pan-genomic islands in the *Hez* genomic repertoire (Fig 2). These islands encode diverse functions and hypothetical proteins with no known function, making comprehensive annotation challenging. The functional description of all the islands, their encoded genes, and adjacent core genes is listed in Table S3. Some islands exhibit preferred geographic distributions corresponding to strains from regionally delimited groups and populations from Spain and Israel (Fig 1). Specific islands, like 3, 9, 13+14, are exclusively found in the genomes of the Israeli clade, indicating divergence and accumulation of rare genes in these regionally restricted groups. Islands 6, 18, 24, 27, 46, and 50 are predominantly found in the genomes of the Spanish strains but occur rarely in strains from other regions. In the case of Islands 15 + 16, which invade between the same core genes, regionally restricted groups (from both Spain and Israel) only have Island 15, while the other genomes spanning multiple regions and populations have Island 16 instead. For several islands we identified putative functions or conserved domains (Islands 4, 5, 7, 15, 18, 24, 31-33, and 44-47) such as: von Willebrand Factor type A domains, intracellular signaling, and sulfur metabolism (see Table S3 for full descriptions). The distribution among isolates suggests that these islands play roles in the specialization and niche adaptation of different *Hez* strains.

### Mobile islands encode ensembles of hypothetical proteins

Notably, 19 islands contained integrases; these islands are denoted in Fig 2 and summarized in Tables S3,4. Most of these islands displayed distinct G+C content (40-55%, all but Islands 21+22) compared to the average of the core genome (~68%). While these islands encode recombinases or integrases, these genes are accompanied by ensembles of hypothetical proteins, for which annotations are unavailable. This is unsurprising as mobile genetic elements encode many unannotated proteins due to their propensity for mobility and fast evolution (32, 33). We used geNomad (34) to identify and classify the viruses and plasmids present in each genome assembly. Only Islands 36 and 38 contained enough hallmarks and marker genes to be classified as plasmids or viruses, both islands were assigned as a member of the tailed double-stranded DNA virus class Caudoviricetes. The absence of viral/plasmid hallmark proteins on the other islands suggests that they may not be actively replicating as a separate entity from the chromosome and represent remnants of integrated mobile genetic elements. The G+C deviation (Fig 2), rarity throughout the pan-genome (Fig 2), and enrichment for uncharacterized genes (Tables S3-4) are attributes these integrase containing islands share.

Six of these islands encoded genes for defense systems against foreign DNA in the form of restriction modification system (RMS) genes (mostly Type I, II systems; see Table S5). Of the 94 RMS-related genes in the entire *Hez* pan-genome, 10 were found on islands (a heatmap summarizing this distribution can be found in Fig S3). RMS encoding islands are rare, each observed in only 2-10 genomes (Table S5). Island-encoded RMS genes exclusively occurred on integrase-containing islands suggesting selection for transmissibility and evasion of resistance mechanisms. Transmissibility is a means to circumvent resistance mechanisms innovated by other mobile genetic elements, that may have been the target of these RMS genes. The Supplemental Text includes a more detailed discussion of RMS systems.

### Transferability of flexible gene content encoded on pan-genomic islands

While many islands have diverged and accumulated genome specific mutations, 11 islands have an identical nucleotide sequence across the genomes they are found in (Fig. 2). In all cases, except Island 38, these identical islands are found in different geographical regions, different clans, and have a limited distribution amongst isolates from the same region. The sequence of Island 11, for an example, is found identically in genomes from Mongolia, Spain, Israel, and Iran. However, the island is still rare in those regions with only 1-2 genomes per location having acquired it. The coexistence of strains with and without the island in the same environment demonstrates the potential utility of that locus. When islands with limited distribution are found across different geographic regions, with an identical sequence, horizontal transfer and migration are plausible explanations for their distribution. We inferred the number of duplications, transfer, and loss (DTL) events for all 50 genomic islands and core genes (2518 total) via topological reconciliation of gene and the species tree (Fig 1) using RangerDTL (35). Genes on a genomic island were concatenated to build an island tree. The genomic islands were inferred as transferred and lost significantly more frequently (two-sided T-test p < 0.01) compared to core genes. After normalization, the average transfer events inferred in the genomic islands was 0.23 (events / (internode x total branch length), while it was 0.15 (events / (internode x total branch length) for core genes. In addition, there were significantly more gene loss events (p < 0.001) in genomic island trees as compared to core genes. The full report of DTL events inferred for every gene tree can be found in the Table S6. While we can infer DTL events for those islands where tree building is possible (i.e., there are at least 3 taxa and the sequences are not identical), the presence of doubleton islands also are likely the result of a transfer if the island is found across multiple geographic regions or phylogenetic groups (e.g., Island 10, 38).

The loci containing these pan-genomic islands are enriched in mobilizing genetic elements. After island discovery, the invasion point (or the immediate two core genes) was expanded to include the genomic neighborhood ±5 genes upstream and downstream of the island. The distribution of tRNAs (T), transposases and integrases in these expanded gene neighborhoods is mapped in Fig 2. In addition, information about transposase families near the islands are included in Table S7. The propensity of tRNA to form hairpin loops paves the way for the integration of mobile genetic elements (36) and horizontal transfer in *Hez*. While there are 70 total tRNAs encoding genes in the genome of *Hez* (37), 18 of them are proximal to these genomic islands. The discovered genomic islands (in all isolates) encode a total for 351 genes, or only 9.5% of ORFs (combined island ORFs from all isolates), while ~25% of all tRNAs in the genome are in the 5 gene neighborhood of an island insertion point, while not being found on the island itself.

### Homeocassettes serve as reservoirs of genetic and gene content variation

Several pan-genomic islands are found between the same core genes, in different genomes, while maintaining local synteny (Fig 2, adjacent purple lines of the same height show same core gene invasion; Table S8 lists these core genes and their predicted functions). The presence of island pairs is mutually exclusive, meaning that, if one genome has island 1, it does not have island 2, except for strain: AMM015 (see Supplemental Text). Additionally, gene sequences encoded on every island are exclusive, meaning they are not recombined from other parts of the genome (although divergent homologs of these genes may exist elsewhere in the genome). These island pairs reflect cassette-level variation in the *Hez* pan-genome. The combined genome representation in every island pair (except islands 13+14 [which is restricted to Israeli strains] and island 44+45), is consistent with the distribution of a core gene (present in all genomes). Six of nine of these island pairs (e.g., island 1 and island 2) are ascribed similar functions and share homologs, but the homologous ORFs on conflicting islands are dissimilar enough (in terms of sequence identity and gene neighborhood) to escape clustering into the homologous gene family on the opposing island (Table 1, Fig 3).

**Figure 3.**
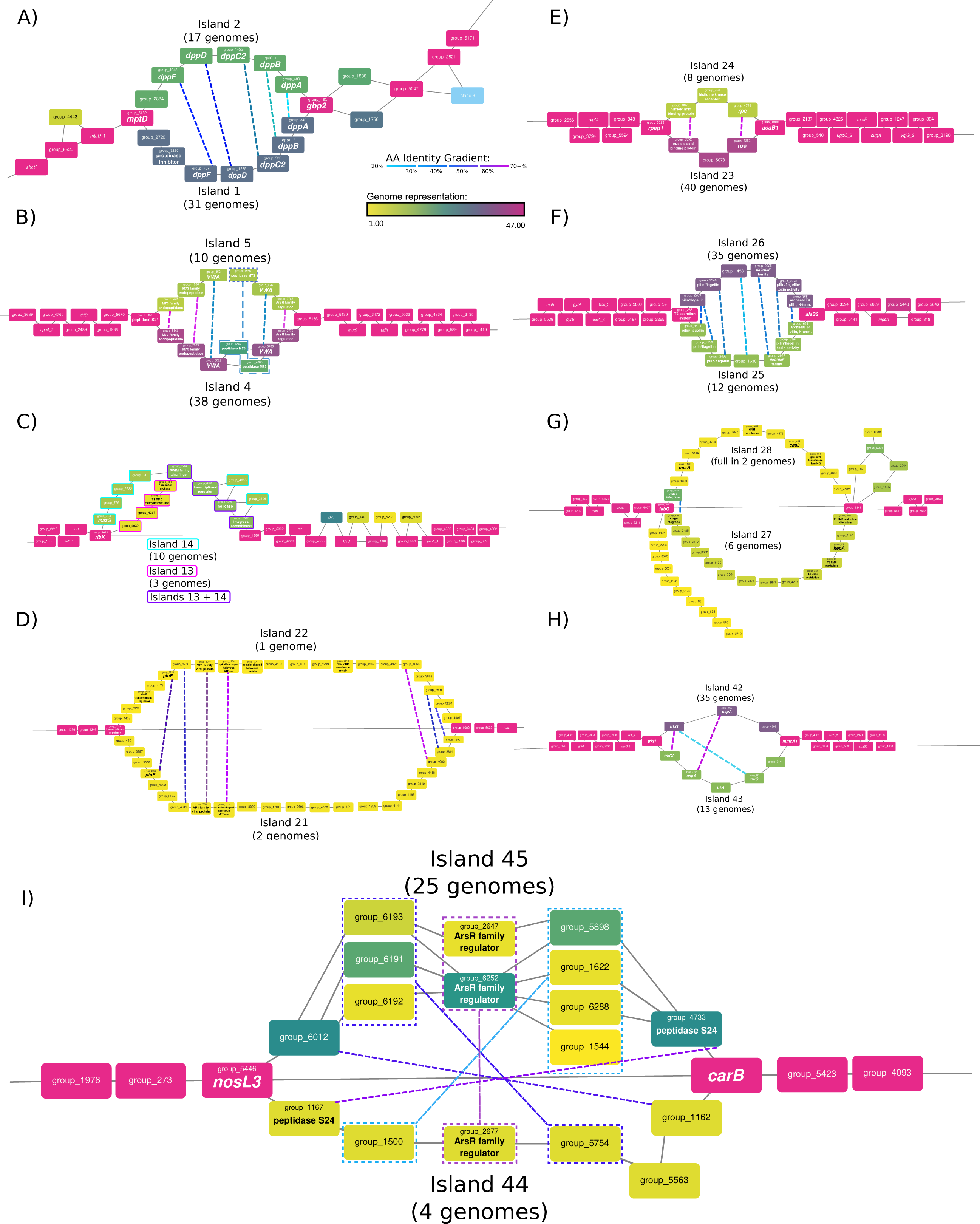
Pan-genome graph depiction of conflicting islands. Close-ups of several regions of interest in the overall pan-genome graph, which show two pan-genomic islands occupying the same locus. Nodes in each graph represent pan-gene families, while edges indicate contiguity between nodes. Core genes are colored magenta and are represented in at least 44 of 47 isolate genomes. Core genes in these depictions are on average 99% identical on the nucleotide level. Islands in between the same core genes are mutually exclusive, and dashed lines between genes on opposing islands indicate homology (coloring indicates % identity). When annotations for pan-genes are available they were written inside the node in the larger font, otherwise arbitrary group_names were assigned. A) Both islands encode the components of a putative oligopeptide ABC transporter, *dppABCDF*. Both islands are inserted between the same core genes (*gbp2* and *mptD*) and display mutual exclusivity barring the AMM015 strain (which increases the total genome number to 48). These two islands are considered a homeocassette. B) These homeocassettes encode for hypothetical proteins which have peptidase and von Willebrand factor type A (VWA) domains, but their specific functions are unknown. Note the slightly different coloration of homeoallele pair group_992 and group_5566, the strains C191 and Hxorientale encode all of the components of Island 5, but have swapped group_992 for its homeallele group_5566 on Island 4. C) Two islands which share genes but do not have any homeoalleles so this pair is not considered a homeocasette. Those genes with the cyan border are exclusive to island 14, while those with the magenta border are exclusive to Island 13; those genes with the purple border are found in both islands. D) These rare islands seem to have indications of viral origins (seen in the shared integrase, Vp1, and haloviral proteins), and have a large G+C content deviation from the core genes. Note the edge which directly connects these two core genes and skips both islands entirely, this indicates that in 44 genomes both islands are not present. E) Two islands inserted between rpap1 and acaB1, sharing two homeoalleles sharing high amino acid identity. F) These islands encode for a Type 4 pilus system and insert between the pilin assembly protein tadC and an alanyl-tRNA ligase. Homologs were found for every encoded gene on opposing islands, and in one case a gene from Island 26 (group_2789) was duplicated on Island 25. G) Two islands of mostly hypothetical proteins, sharing only an integrase integrated between fabG and group_5345. Both islands encode functions related to defense (homing endonuclease and RMS genes). The two anchoring core genes show evidence of integration with other elements that are not contiguous with the core genome, indicating that these genes may be a hotspot for integration or recombination. Note the edge which connects the two anchoring core genes, which displays contiguity of those core genes in the absence of the two islands. The loop (and those like in the the pan-genome graph) extending from group_5345 (and includes group_182, group_6371, group_6008, group_2044, group_1055 and) ends again back at group_5345 is not considered an island in this work. These genes are not a contiguous set that is found in multiple genomes and only have one core gene connection, and are actually the result of multiple paths between three dead ends. These three paths are: 1) groups 5345-182-6371-6008 (shared in 3 genomes), 2) groups 5345-1055-2044-6371 (shared in 10 genomes), and 3) group_6371 (1 genome only). An edge is drawn between any nodes (group_6371 is contiguous with all three paths) which are found to be contiguous in any of the genomes (even if it is only present in 1 genome), this is reflected in the difference in coloring of the nodes in the loop. H) Two islands which encode components related to potassium import, integrated between the core genes trkH and mmcA1. The two islands share 3 homeoalleles (pairs are indicated with dashed lines), where there is an unbalanced duplication of the trkG (membrane component of trk transporter; also called trkH) gene on Island 43. In addition, Island 43 has another copy of the trkA transporter (albeit truncated), where a core gene version of this gene exists upstream of the islands. I) These islands insert between the core genes *nosL3* and *carB,* and encode an ensemble of mostly hypothetical proteins. The orientation of the opposing islands is reversed in this example, and additionally, island 45 has underwent numerous reshuffling and reorganization events. Island 45 can be seen as 3 different islands (i.e., it has at least 3 different paths to reach the next core gene). Although the islands have been reorganized genes on opposing islands are homologous with various degrees of amino acid identity. Genes that have been duplicated (indicated by the dashed box) are homologous.

**Table 1.**
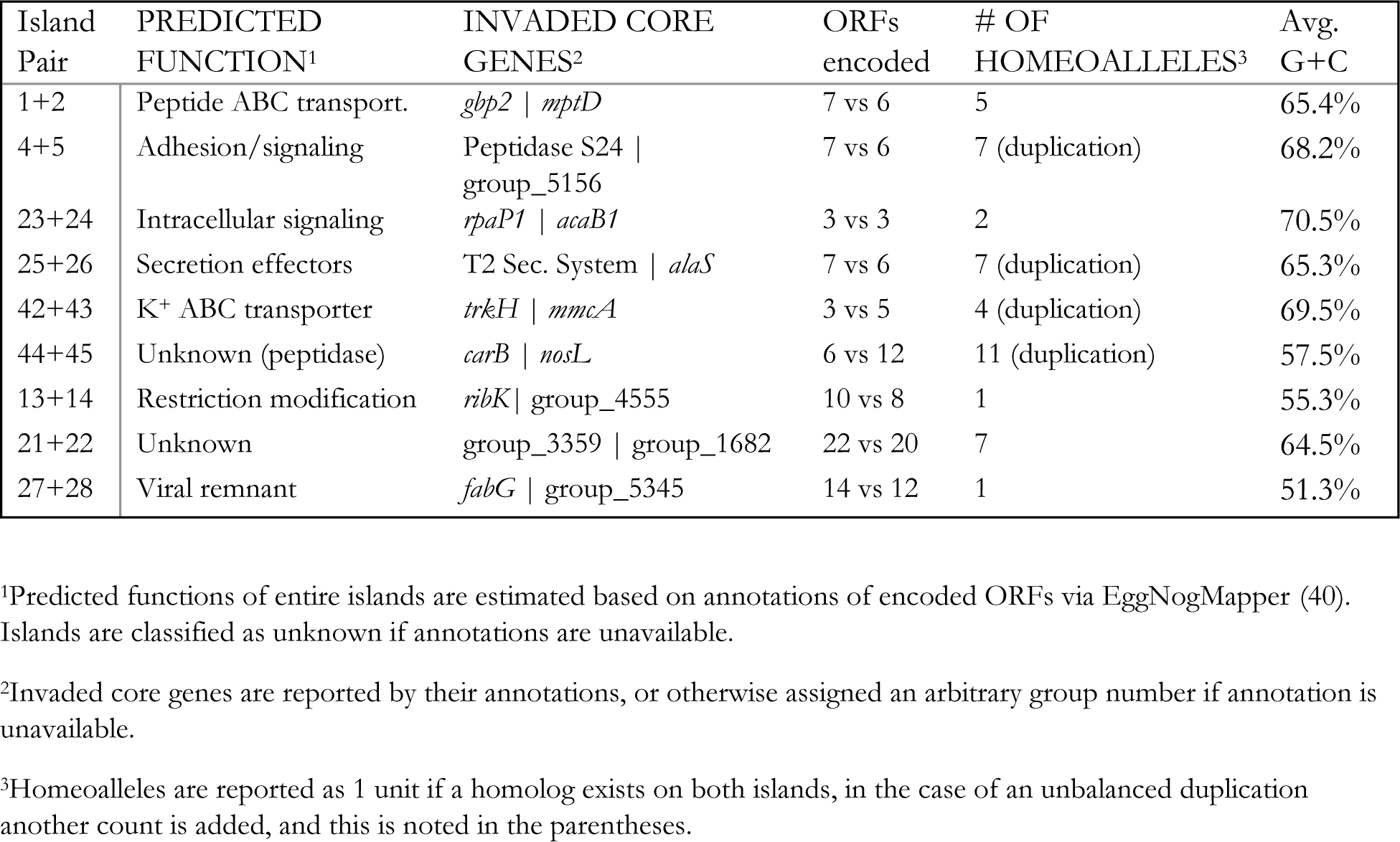
Summary of Conflicting Island Pairs.

| Island Pair | PREDICTED FUNCTION<sup>1</sup> | INVADED CORE GENES<sup>2</sup> | ORFs encoded | # OF HOMEOALLELES<sup>3</sup> | Avg. G+C |
| --- | --- | --- | --- | --- | --- |
| 1+2 | Peptide ABC transport. | *gbp2* \| *mptD* | 7 vs 6 | 5 | 65.4% |
| 4+5 | Adhesion/signaling | Peptidase S24 \| group_5156 | 7 vs 6 | 7 (duplication) | 68.2% |
| 23+24 | Intracellular signaling | *rpaP1* \| *acaB1* | 3 vs 3 | 2 | 70.5% |
| 25+26 | Secretion effectors | T2 Sec. System \| *alaS* | 7 vs 6 | 7 (duplication) | 65.3% |
| 42+43 | K<sup>+</sup> ABC transporter | *trkH* \| *mmcA* | 3 vs 5 | 4 (duplication) | 69.5% |
| 44+45 | Unknown (peptidase) | *carB* \| *nosL* | 6 vs 12 | 11 (duplication) | 57.5% |
| 13+14 | Restriction modification | *ribK* \| group_4555 | 10 vs 8 | 1 | 55.3% |
| 21+22 | Unknown | group_3359 \| group_1682 | 22 vs 20 | 7 | 64.5% |
| 27+28 | Viral remnant | *fabG* \| group_5345 | 14 vs 12 | 1 | 51.3% |

<sup>1</sup>Predicted functions of entire islands are estimated based on annotations of encoded ORFs via EggNogMapper (40). Islands are classified as unknown if annotations are unavailable.
<sup>2</sup>Invasion core genes are reported by their annotations, or otherwise assigned an arbitrary group number if annotation is unavailable.
<sup>3</sup>Homeoalleles are reported as 1 unit if a homolog exists on both islands, in the case of an unbalanced duplication another count is added, and this is noted in the parentheses.

We considered every pairwise combination of genes on each conflicting island pair (where there was not a large difference in the number of ORFs encoded within the set) with blastn, blastp, and PRSS searches (Table S8). Genes on one island which have homolog(s) on the opposing island were termed “homeoalleles”, while genes that do not have homologs on an opposing island were termed “iORFans” (as in “island encoded orphan gene”). Homeoalleles, not to be confused with homeologs in eukaryotes (i.e., a homolog resulting from allopolyploidy), are divergent iso-functional variations of a gene that may be swapped/transferred within an exchange group often in equivalent genomic positions (38, 39). In Islands 4+5, we observed an instance of within island recombination where a peptidase from Island 5 was swapped with its homeoallele on Island 4 in the strains C191 and Hxorientale (Fig 3b), while the other 10 strains which have Island 5 retained the original peptidase. Thus, we term conflicting genomic islands which share homeoalleles as “homeocassettes”. We defined homeocassettes as any two conflicting islands which have at least 50% of their gene content as homeoalleles. This somewhat arbitrary threshold allows the distinction between conflicting islands which share many homeoalleles (Fig 3ab) and those that are likely viral remnants which only share the integrase gene (Fig 3c).

In total, we identified 9 conflicting island pairs (18 islands total; Fig 3, Table 1). Six island pairs were enriched in homeoalleles, forming homeocassettes, while three pairs had more iORFans and shared at least an integrase/recombinase with the opposing island. The genes in the islands of a homeocasette pair (Islands 1+2, 4+5, 23+24, 25+26, 42+43, 44+45; Table 1) are ascribed similar or identical functions. In contrast, the conflicting island pairs Islands 13+14, 21+22, and 27+28 do not share similar functions and these loci appear to be hotspots for recombination (Table 1, Table S3). Opposing islands of a homeocassette were found across geographic locations and phylogenetic groups (Fig 2). For example, Israeli and Spanish strains have a mixture of island 1 and island 2 within each geographic location.

Homeocassettes encode diverse functions, and each island within a pair offers a different variation upon that function. They also display exclusive genetic variation and selective organization of gene cassettes/operons for a set of homologs. The observation that homeocassette pairs are located between the same core genes, appears to reflect their integration through homologous recombination in the conserved flanking region following genetic transfer or mating (41). The homeocassette pairs we observed in the *Hez* genomes differ in the number of encoded ORFs (Fig 3, Table 1); this difference is due to iORFans or paralogues found on either island. The presence of multiple, presumably synergistic genes or operons on the islands allows the host to gain entire functions with the acquisition of an island. Due to the amino acid sequence divergence displayed by homeocassettes it may be improper to consider them iso-functional *in toto*. Similar to homeoalleles of aminoacyl tRNA synthetases (42), the main function of a homeoallele pair may be the same; however, it is likely that the pair differs in some respects that impact organismal fitness under some environmental conditions, which would ultimately affect the persistence and spread of the islands within the population.

The retention of homologous, but highly divergent GIs in equivalent genome positions may facilitate the adaptation to specific environmental niches. The islands might encode weakly selected functions *sensu* Lawrence and Roth (Lawrence and Roth, 1996), and this would have contributed to their faster divergence compared to the core genes. The replacement or divergence of loci between otherwise closely related strains of the same species has been previously observed in *Halobacterium salinarum* (43) and *Haloquadratum walsbyii* (44), which is referred to in those works as “divergent segments (DivSEG)” or “deletion-coupled insertion”. *H. salinarum* strains 91-R6 and R1 differ in at least two loci (DivSEG04 and DivSEG18) in equivalent genome positions which encode homeoalleles that share up to 80% AA identity (between their respective genomes; (43)). *Halquadratum walsbyi* strains C23 and HBSQ001 have a divergent region (in terms of sequence and length) where an S-layer glycoprotein in the region only shares 55% AA identity, while the surrounding genes on average exhibit 97% sequence identity (44). Additionally, the Cas effector modules, in a wide sampling of organisms, have been observed to have been swapped with divergent homologs trafficked by mobile genetic elements (45) These modules insert adjacent to the core components of Cas4, Cas1 and Cas2, which remain in place, retain synteny and relatively high sequence identity. In these cases and the homeocassettes describe in this study, the divergence between homeoalleles is greater than the divergence observed between the core genes. Fig 3 demonstrates the amino acid divergence between genes of an island pair (from 35%-70%), while the nearest neighboring core gene is ~99% similar. The contrast in divergence in colocalized genes may be the consequence of HGT or an acceleration of evolutionary rate for specific reasons. The prevalence of similar replacement and divergence dynamics outside of *H. ezzemoulense* suggests that this may be a biological feature that has a wider distribution in haloarchaea.

Different variants of the same gene cassette may offer unique advantages in different conditions, especially if the environment is heterogenous. Those islands which encode surface components (such as Islands 1+2, 4+5; see Table S3) can serve as viral attachment receptors. Multiple viral receptors in a single meta-population may confer immunity to predators in a sub-population. The co-occurrence of homeocassette pairs in the same region (in different genomes) and the swap of endopeptidase homeoalleles observed in Islands 4+5 (in strains C191 and Hxorientale; Fig 3b), indicates that it is possible for an organism to switch between the contents of a pair through recombination of the conserved surrounding regions. It is also possible that homeocasettes may play a role in speciation by acting as the crystallization points for expanding regions of divergence where barriers to within group recombination are erected. This assertion may be at odds with the HGT and genome dynamics known for the Haloarchaea (41). However, the divergence found between homeocassettes is similar to the loci of differential divergence described in *Escherichia* and *Salmonella* (Retchless & Lawrence, 2007), whereby local genetic isolation was maintained for tens of millions of years before the divergence of the two lineages.

### Differential solute transporter utilization amongst *Hez* isolates

10 islands encode the components for various transporters: 8 ABC transporters and 2 trk-type transporters (Table S9). These transporter islands (Table S9) have an average G+C which closely resembles the *Hez* core genome. In contrast to islands which encode integrases, almost every gene on transporter islands could be annotated. Furthermore, transporter islands have high genome representation, suggesting the genes encoded on them have greater utility and therefore higher lineage persistence than genes found on integrase encoding islands. Transporter encoding islands illustrate the utility of the pan-genome graph for GI discovery, as they do not display obvious organizational deviations from the genome average. These islands are likely not the result of HGT from outside *Hez*, and seem to have evolved, recombined, and spread within *Hez* and earlier related lineages. Phylogenies of phosphate and peptide transporter islands were inferred using the sequences of the more conserved ATP binding subunits (Fig S4,5), and included a sampling of haloarchaea outside of *Hez.* These trees reveal that these islands existed in the earlier haloarchaeal lineages (i.e., before the divergence of *Hez*) but also have a limited distribution (i.e., not every member of the same genus/species encode the island uniformly). The limited distribution displayed by transporter islands reflects differential availability or utilization of environmental solutes amongst members of *Hez*, otherwise the transporters would be completely fixed or lost in the population. These discovered transporters have putatively ascribed substrates which include reduced carbon compounds, phosphate, polyamines, small peptides and potassium ions. The physiological and evolutionary implications of these transporter islands are discussed in extensive detail in the Supplementary Text.

Island 19 is the sole instance of a complete carbohydrate uptake transporter (CUT) in the *Hez* pan-genome, found in a few Iranian, Mongolian, and Spanish strains (6 total), but notably absent in Israeli isolates. The presence of homologs to genes from the two described CUT operons from *Pyrococcus furiosus* (46), suggests its ability to import both trehalose/maltose and maltodextrin. Strains which harbor this island have a greater range of nutritional competence when those carbohydrates are present in the environment.

Polyamines (such as agmatin) reportedly play crucial roles in Archaea, modifying translation factors and tRNA, which are essential for translation (47–49). Earlier studies noted the complete absence of polyamine biosynthesis genes in haloarchaea (50). Later studies reported the discovery of agmatinase (*speB*) but noted the lack of a polyamine import mechanism for which *speB* would act in concert (49). Island 16 is a sporadic island which encodes 5 genes for polyamine import (*potABCD* system (51)). This presents a possible polyamine scavenging system which confers a growth advantage for those strains which have acquired Island 16 (Fig 2) in environments where polyamines are present.

Islands 42+43 in *Hez* are homeocassettes that encode TrkAH transporters (Fig 3h). Unlike other transporter islands, these TrkAH transporters are not energized by ATP hydrolysis (52). K+ import plays a critical role for the salt-in strategy employed by *Hez* and other haloarchaea (53). Island 43 stands out as it contains a variant of the *trkA* core gene found upstream of the homeocasette insertion point (Fig 3h). The core TrkA has 480 amino acids, while the island-encoded TrkA is shorter, consisting of 230 amino acids and displays high similarity to the N-terminal portion of the core TrkA. Previously, a similar truncated *trkA* gene in *Thermotoga maritima* was considered a pseudogene due to a frameshift mutation (54, 55). However, complementation assays (Johnson et al., 2009) revealed that these truncated TrkA proteins, along with a novel membrane-spanning protein, form a new sub-class of K+ importers known as the two-subunit TrkA importer (more in Supplemental Text). The existence of Islands 42+43, and the core TrkAH genes highlights the diverse potassium import strategies present in the *Hez* pan-genome (Fig 3h).

5 of the transporter islands encode divergent (up to 50% amino acid identity) putative di/oligopeptide transporters (*dppABCDF* operon), including two homeocasette pairs (Fig 3a). The core genes of these 5 islands include the necessary components of an ABC transporter which traffics small peptides, however, the ATP coupling proteins DppD+F are fused into one ORF on Island 12 (Fig S4). Accessory genes on these islands (i.e., those genes not homologous to the ABC transporter components) are involved in amino acid metabolism, suggesting these islands genuinely import peptides. Strains with Island 39 demonstrate uncertainty in its classification as a plasmid or integrated into the chromosome. These findings are consistent with the notion that pan-genomic islands act as reservoirs of genetic variation, allowing the retention of divergent gene cassettes over time. These differences between the islands suggest each island has diverged to accommodate slightly different peptide substrates. Consequently, different strains of *Hez* can adapt to different peptide availabilities in different environments and have a greater range of niche adaptability. Genomes which have multiple versions of *dpp* transporters (such as UG_123 and UG_126 encode islands 2 and 41; for more examples see Fig 2) may benefit from a dosage effect where they are able to extract more peptides from the environment by encoding more transporters.

### Island encoded phosphate transporter is a variation of a homologous core gene neighborhood

Island 49 encodes a variant of the *phnCDE* operon, which was experimentally associated with eDNA uptake in conditions where phosphorous was a limiting nutrient in *Haloferax volcanii* (56). This GI was found in 15 genomes across different geographic regions. A homologous, but divergent *phnCDE* gene cluster was also found in the *Hez* core genome over 2.36M bp downstream of the island insertion point (Fig S5). The *Hez* core-*phnCDE* cluster displays greater similarity to the *Haloferax volcanii* phnCDE system in which the knockout experiments were performed (see similarity comparison is Table S8). Island 49 encodes not only the *phnCDE* homologs, but also contains a duplicated permease (*phnE2*) and a HAD hydrolase which affirms the island’s association with phosphate/phosphonate import (see the Supplemental Text). The island encoded *phnE1* and *phnE2* genes are also present in *Haloferax volcanii,* where they are encoded on the megaplasmid pHV4, but this gene neighborhood does not contain homologs of the *phnCD* genes (Fig S5).

The presence of two homologous phosphate uptake systems in *Hez* demonstrates the role of pan-genomic islands in diversity generation. Many of these islands and homeocassettes contain the result of an unbalanced (i.e., uneven ratio of homologs on opposing islands, such as 2 phnEs vs 1 phnE) gene duplication (Table S8, the duplications are highlighted in yellow). Many of these duplications are not ancestral, as these islands have a sporadic distribution across regions and phylogenetic groups (Fig 2). In this work, duplicated genes tend to reassociate with homologous, but divergent variants of genes possibly recombined from the source gene neighborhood, this is seen in several of the illustrated examples (Island 49: Fig S5, Islands 4+5: Fig 3b, Islands 25+26: Fig 3f). The fate of duplicated genes is often pseudogenization (57), and in some cases neofunctionalization In the case of *phnCDE* system on Island 49 the later seems to be the case. This is because homologs of the *phnCD* genes are also found on the same island. While the acquisition or duplication of a core gene may seem wasteful and counter-productive to fitness (redundant genes and energetic costs), it may be the starting point of genetic divergence or lead to greater niche adaptability in the species. Retchless and Lawrence (58) showed that the acquisition of an adaptive gene can be the starting point of genome divergence spreading out from this gene into the gene’s neighborhood. This acquisition event is not dissimilar to a gene duplication into a distant locus, as seen in Island 49. The reconstitution of homologous genes from the source gene neighborhood in the duplicated gene neighborhood illustrates the selective/selfish manner with which gene neighborhoods form. We speculate that the duplicated gene has a greater chance of persistence and replication if it can be transferred and utilized in synergy with the other homologous components of the same system.

## Conclusion

Using a pan-genomic framework, namely the pan-genome graph, for the discovery of islands has a few advantages. Principally, this method allows for the detection of islands that have the same underlying features (GC content, gene density, etc.) as the genome of the host. Discovery of genomic islands using the pan-genome graph does not rely on a reference genome and is extensible to any dataset containing highly related (and syntenic) genomes. Homeocassettes and those islands encoding ABC transporters would not be detected with a method that focuses on deviations from features in the core genome. Additionally, a sense of reliability and support can be attributed to islands found under this framework. All the islands must be syntenic, or the graph pathing will fork and create singleton islands that would corrupt or complicate (Fig. 3g,h) the order of genes. This graph-based framework allows for the exploration of gene neighborhood surrounding the island, in all the included genomes, which can uncover mobilizing elements that may explain the ontology of the island.

Broadly, there seems to be few barriers to horizontal transfer between strains of *Halorubrum ezzemoulense* in distant geographic locales, although strains in the regionally delimited groups (Fig 1; highlighted clades) share more GIs with each other than with those strains outside the groups. Pan-genomic islands play an important role in the generation and upkeep of within species variability and persist as divergent units even if two islands occupy the same locus. The discovered transporter islands possibly provide a greater range of niche adaptability, while not changing the fundamental genetics of a species member. Rare island encoded genes are possibly made accessible to other species members via recombination using the conserved surrounding regions as a template. Some genomic islands, however, harbor or are the remnants of selfish genetic elements, which take advantage of *Hez’s* propensity for recombination to persist and spread to new uninvaded populations. These findings illustrate how islands confer genetic and gene content variation to *Hez*.

## Methods

### Isolates collection, sequencing, and genome assembly

Spanish isolates were collected and their purified DNA sequenced using the methods described in (Feng et al., 2019). Israeli strains were isolated using themethods described in (59): water samples from the Dead Sea (DSW) were extracted using Niskin bottles and diluted with autoclaved double-distilled (DDW) water (1/5 [DDW/DSW] ratio. The mixture was amended with 0.1% glycerol, 1µM K_2_HPO_4_, 1g/l peptone, 1g/l casamino acids and incubated at 30° for 42 days. DNA for both types of samples were purified using the DNeasy PowerLyzer PowerSoil kit (using the manufacturer’s protocol). DNA was purified from the 0.22µM filters and used for library preparation and ran on Illumina NovaSeq with SP flow cell, generating paired end reads (2 x 150 bp). Reads were assembled with SPAdesv3.11.1, on default parameters using 150 bp reads. The genome of *Hez* strain fb21 was assembled using a combination of short and long Pac-Bio reads, as described in Feng et al., 2019. Genomes were also mined from NCBI to supplement the in-house isolates; their accession numbers are provided in Table S2. Genome assembly quality was assessed with CheckMv1.0.7 (60), using the 149 Archaeal marker set. Genomes with low completeness and high contamination were discarded, the genomes quality report is shown in Table S2. In addition, the qualities of the genomes were processed by the panaroo-qc script (27), and additionaloutlying genomes were removed.

### Genome distance calculation and tANI phylogeny

Genome distance was calculated using an extension of the jANI algorithm (Gosselin et al., 2021.), which also takes into account the alignment fraction (AF) of the genomes. The genome distances were used to construct a phylogenetic tree, with bootstrap support per the tANI method. Genomes with a distance <0.315 (with AF =1, this corresponds to ANI >73%) are considered members of the same species. The complete distance matrix is displayed in Table S1, the resulting phylogeny is depicted in Figure 1.

### Gene calling, gene family clustering, and pan-genome graph generation

ORFs were predicted and annotated in these genomes using Prokkav1.14 (61), under the same gene model (trained for *Hez* genomes), under the default settings and with the Archaea flag. In addition, all gene families were cross annotated with EggNOG-Mapper5 (40), and several genes of interest were analyzed in in HHPred (62). Predicted ORFs were clustered into gene families using multiple steps: 1) identity threshold (98%), and 2) graph-based-clustering where gene families from different genomes are collapsed into the same family if their local synteny is conserved throughout multiple corrections and filtering of the graph, as described in Tonkin-Hill, et al., 2020 (27). The resulting clustered gene family nucleotide sequences were aligned with MAFFT (Atoh and Standley, 2013), and their distributions can be visualized in the pan-genome matrix. The pan-genome graph expresses every gene family and how they are ordered across all genomes, close-up examples of the graph are contained in Fig 3. The interactive graph is found in File S1 and can be visualized in Cytoscape (64).

### Querying the pan-genome graph for genomic islands

The pan-genome graph contains every single gene family in the pangenome (as nodes) and shows how each gene connects to the next (across all genomes) with its edges (which can be multifurcating). The graph itself was generated with the Panaroo pipeline, using –clean-mode strict and default parameters (27). The graph was traversed using the accessory script panaroo-gene-neighborhood, with –expand_no 20 and default parameters. This script was used for calculating the gene neighborhood paths for all 8,401 pan-genes, this provided the ±20 gene neighborhood for each gene. In this dataset, core genes have a representation of 44+ (in all genomes) and cloud genes have a representation in only one genome. The representation of all gene families (i.e., genome coverage or how many genomes a pan-gene is in) was overlayed onto the previously calculated gene neighborhood paths.

Genomic islands, by definition, cannot contain a core gene; but are often found invading between two core genes. These genomic islands always adhere to a simple representation pattern in this pan-genomic framework: core gene-(rare gene(s))-core gene. We searched the ±20 gene neighborhood paths (which now includes the representation of each gene) for the following coverage (representation) pattern: “44+”-“(1–43)x”-“44+”, where x is ≥2. This returned any multi-gene fragment, of low coverage, that is contiguous with two flanking core genes (high coverage). The flanking core genes were retrieved for each insertion, and the adjacent island was visually inspected in the pangenome graph (final_graph.gml; File S1) in Cytoscape (64). Only islands which contain few multifurcations and have a traceable path from one flanking core gene to the other was reported (50 total). Islands with complicated multifurcations, multiple core gene invasion, and divergent synteny were discarded (131 total). This approach provides a measure of reliability (i.e., coverage) by stipulating: 1) the inclusion of the invaded core genes (which has at least 95% coverage), and 2) the local synteny of the genomic island is intact (across all genomes which include the island). A hard-coded script of this search process can be found in the supplemental material as an interactive Jupyter Notebook (File S2).

### Analysis of *Hez* population structure

The Tajima’s D (65) statistic was calculated for every core gene (i.e., a gene that is found in at least 44 of the 47 genomes) alignment using Pegas (66), those gene families with a Tajima’s D <|0.2| were considered as not seen by purifying selection (i.e., almost neutral). Neutral core genes were concatenated and used to generate a SNP matrix. These genomes were hierarchically clustered, based on SNP distances, into populations using RhierBAPs (29). The population membership for every genome was then mapped back to the tANI tree (Figure 1).

### Inference of gene transfers

Proto-phylogenies (gene trees without correction applied) were calculated for all genes of interests and concatenates (islands). Proto-phylogenies, the associated nucleotide alignment, and the species tree (Figure 1) were supplied to treefixDTL (67) to produce the corrected phylogeny where HGT, topological information, and sequence likelihoods (probability of observing the sequences estimated using a pairwise distance matrix of the alignment under the JTT model) were considered. The corrected phylogenies were optimally rooted based on minimizing DTL reconciliation costs in OptRoot. These rooted phylogenies and the species tree were then supplied to RangerDTL2.0 (35) where duplication, transfer, and loss events were inferred. Each gene tree in this analysis was explored through 300 independent runs in the reconciliation space using 3 different DTL costs ([2,3,1], [3,3,1], [2,4,1], costs for a [duplication, transfer, loss]). Inferred DTL events for each gene tree were normalized using the number of internal nodes and total branch length of the gene tree.

### Identification of Restriction Modification Systems Genes

Query sequences for RMS genes were gathered using a slight modification of the method described by Fullmer et al., 2019. Briefly, all sequences on REBase were downloaded, and clustered using a 70% identity threshold in USearch. HMMs were built for all clusters and used to search the clustered gene families of the *Hez* pan-genome. Those gene families that matched an RMS gene, had their genomic neighborhood (5 genes upstream/downstream) searched for other RMS genes. RMS gene type assignment (Type 1-4) was determined by the best ReBase hit, and in the case of ambiguity for multiple RMS genes in the same neighborhood the assignment was typed using the restriction enzyme’s annotation.

## Author Contributions

The project was conceived by YF, ASL, and JPG. Sampling, sequencing, and genome assembly of the Israeli samples was conducted by N.A-P. and U.G., while A.S.L. and R.T.P was responsible for the Spanish and Iranian samples. D.A. and Y.F. performed the pan-genomic analyses. All authors contributed to the writing and editing of the manuscript.

## Data Availability

New *Hez* draft genomes from Israel are found under BioProject PRJNA925523, and those genomes from Spain are found under BioProject PRJNA925515. Additionally, genome gff files, nucleotide sequences of island encoded genes, the pan-genome graph and additional supporting materials are deposited in supplemental_hezpg.zip.

## Supporting information

All of the supplemental material

## Acknowledgements

This work was supported by grants from the Binational Science Foundation [BSF 2013061 to U.G., J.P.G., and R.T.P.]; the National Science Foundation [NSF/MCB 1716046 to U.G., R.T.P., and J.P.G.]. We thank Dr. Andrea Makkay for sequencing the Spanish strains. We also thank Dr. Anna Green (University of Massachusetts, Amherst) for her insightful discussions of pan-genomics.

## Supplemental Material Table of Contents

### Figures

Figure S1. **A description of the *Halorubrum ezzemoulense* pan-genome calculated by Panaroo.**

Figure S2. **2,621 core genes phylogeny of *Hez* isolates.**

Figure S3. **RMS distribution mapped onto phylogeny**.

Figure S4. **Variations of the *dppDF* transporters contained on the islands of *Hez*.**

Figure S5. **Comparison of phnCDE genes in Halorubrum ezzemoulense and Haloferax volcanii.**

### Tables

Table S1. **Information of assembled genomes from isolates.**

Table S2. **Pairwise tANI distance matrix.**

Table S3. **Description of Gene Content Encoded on Islands.**

Table S4. **Information on integrase containing islands.**

Table S5. **Reports of island encoded restriction modification system genes.**

Table S6. **Inferred DTL events for islands and core genes.**

Table S7. **Description of transposable elements found within the proximity of islands.**

Table S8. **Homeoallele searches between opposing islands.**

Table S9. **Descriptions of transporter encoding islands.**

Table S10. **Investigation of AMM015’s competitive genomic islands.**

### Supplemental Text

Restriction modification systems

AMM015 – a strain that harbors both islands of a conflicting pair

The only *Hez* CUT transporter is encoded on a rare island

Island encoded transporters reveal a polyamine scavenging mechanism

Variations of Trk-type K+ import encoded on a homeocassette

Divergent di/oligopeptide transporters are retained in the *Hez* pan-genome on islands

The island encoded phosphate transporter is a variation of a homologous core gene neighborhood

### Supplemental Files

File S1. **Pan-genome graph.** final_graph.gml

Can be accessed through Cytoscape (64).

File S2. **Jupyter notebook of island search process.** find_islands.ipynb

Can be accessed using python or Jupyter notebook affiliated programs.

File S3. **Island gene alignments.** Island_gene_seqs.zip

