## Supplementary material for "Using the pan-genomic framework for the discovery of genomic islands in the haloarchaeon *Halorubrum ezzemoulense*": All of the supplemental material: FileS2.find_islands.ipynb

```
# %% [code]
import glob

# %% [markdown]
## Set up

1. A directory with all gene families in the pan-genome (can be generated with -a pan parameter in Panaroo main pipeline)
2. A directory with all the gene neighborhood paths (can be generated with panaroo-gene-neighbourhood --gene all_genes --graph final_graph.gml -expand_no 20 --out all_genes.walk.txt)

# %% [markdown]
### Parameters
Fill in these values to adjust for your data set:


# %% [code]
number_of_genomes = 0 #change to number of genomes (integer)
core_threshold = 0 # percentage of genomes in which a gene must be found to be considered a core gene (default= 0.95)

min_island_size = 0 #smallest island (in ORFs) to be considered (default = 2)


working_core = (number_of_genomes*core_threshold)

# %% [markdown]
### In the gene families directory:
Calculate the coverage of every gene family, this is saved in the clust_distro dictionary.

# %% [code]
clust_distro ={}
for file in glob.glob("*.fas"):
    with open(file,'r') as ofh:
        seqs = 0
        for line in ofh:
            line = line.strip()
            if line.startswith('>'): #counts the number of sequences
                seqs += 1
    clust_distro[file[:-8]] = seqs

# %% [markdown]
### In the gene neighborhood paths directory:
Grab every gene path and save it into file_paths dictionary

# %% [code]
file_paths = {}
for nayb in glob.glob("*.walk.txt"):
    with open(nayb, 'r') as ofh:
        
        line_num = 0
        for line in ofh:
    
            line = line.strip()
            if not line:
                continue
            if "support" in line:  #skips header
                continue
            
            nayb_line = line.split("\t")[2].split(",")  #retrieves pathing information
            
        
        
            file_paths[nayb[:-9]+ "|" + str(line_num)] = nayb_line
            line_num += 1

# %% [markdown]
### Overlay the gene family coverage numbers onto the gene neighborhood paths:
This uses the clust_distro dictionary to report the coverage numbers onto the gene neighborhood paths. <br>

At this point you can hit run all to execute all blocks.

# %% [code]
file_lines = {}
for nayb in glob.glob("*.walk.txt"):
    with open(nayb, 'r') as ofh:
        
        line_num = 0
        for line in ofh:
            nayb_line =[]
            line = line.strip()
            if not line:
                continue
            if "support" in line:
                continue
            for numberz in line.split("\t")[2].split(","):
                if numberz in clust_distro.keys():
                    nayb_line.append((clust_distro[numberz]))
                else:
                    nayb_line.append((1))
        
        
            file_lines[nayb[:-9]+ "|" + str(line_num)] = nayb_line
            line_num += 1

# %% [markdown]
### Function to search all gene neighborhood coverage patterns


# %% [code]
def check2(coverage_path):
    first_border = 0
    i_hold = 0
    for i,j in enumerate(coverage_path):
        
         
        if j <= (working_core - 1): #skips non-core genes until a core gene is found
            
            continue
        if j > working_core:  ##finds the instance of the first core gene encountered
            if i < len(coverage_path)-1:
                if coverage_path[i+1] >= working_core:
                    continue
                else:
                    i_hold = i +1
                    break
            else:
                break
    for i,j in enumerate(coverage_path[i_hold:]):
        if j >= working_core:
            i_end = i+i_hold
            return(coverage_path, i_hold, i_end)
         
#if all are core genes on the path, returns the path, but ihold would be 0.
#if island pattern is present, returns the path, ihold and iend have contain the slices for the insertion.
#if island pattern is not present, returns nothing. 
                

# %% [markdown]
### Use the function to search all of the gene neighborhood coverage patterns
This prints out a file "patterned_lines.txt" which contains all of the possible islands. Column 1 is the name of the gene in which the gene neighborhood was calculated, column 2 coverage along the gene neighborhood, column 3 and 4 are the indices in that gene neighborhood for the island (which does not include the flanking core genes)

# %% [code]
for keys, vals in file_lines.items():
    ell = check2(vals)
    if ell != None:
        if int(ell[2]) - int(ell[1]) > min_island_size: # only islands of size 2 or more were kept
            print('{} \t {} \t {} \t {}'.format(keys, ell[0], ell[1], ell[2]), file=open("patterned_lines.txt",'a'))
    

# %% [markdown]
### Dereplication
Since the gene neighborhood pathing was done for every single gene family, there will be redundant/overlapping paths (i.e., genes on the same island have similar paths). The following gets rid of these redundancies.

# %% [code]
##reads island information, including the name of the gene and the length of the island and its indices

pattern_check = {} 
with open("patterned_lines.txt",'r') as ofh:
    for line in ofh:
        stored = []
        line = line.strip()
        label = line.split("\t")[0]
   
        stored.append(int(line.split("\t")[2]))
        stored.append(int(line.split("\t")[3]))
        
        
        pattern_check[label] = stored

# %% [code]
actual_pattern = {} #everything in this exhibits the core-rare-core pattern with min length of 2
for kays, veals in pattern_check.items():
    if veals[0] > 0: #this ensures that the island is not at the beginning or end of the gene neighborhood
        actual_pattern[kays] = veals
        
insert_len_underep = {} #checks the length of islands that have invaded the same core genes
for gene_sub, slices in actual_pattern.items():
    insert_len_underep[gene_sub] = (slices[1] - slices[0])
    
    
uniques = [] #finds unique core genes that are invaded
for items in insert_len_underep.keys():
    uniques.append(items.split("|")[0])

# %% [code]
#this finds the longest possible stretch of an island within its core gene insertion, and gets rid of redundant entries
derepped = {}
for jeans, lens in insert_len_underep.items():
    if jeans.split("|")[0] in derepped.keys():
        if lens > derepped[jeans.split("|")[0]]:
            derepped[jeans.split("|")[0]] = lens
        else: 
            continue
    else:
        derepped[jeans.split("|")[0]] = lens
        
        
derepped2 = {}
pop_out = []
for jeans, lens in insert_len_underep.items():
    if lens == derepped[jeans.split("|")[0]]:
        if jeans.split("|")[0] not in pop_out:
            derepped2[jeans.strip()] = lens
            pop_out.append(jeans.split("|")[0])
    else: 
        continue
        
prime_paths = {}
for interest in derepped2.keys():
    slice_check = []
    for segs in file_paths[interest]:
        if segs in derepped.keys():
            slice_check.append(derepped[segs])
        else:
            slice_check.append(0)
    max_indy = np.argmax(slice_check)
    prime_paths[interest]= file_paths[interest][max_indy]
    
primera = []
for primus in set(prime_paths.values()):
    for name in prime_paths.keys():
        if primus == name.split("|")[0]:
            primera.append(name)

# %% [markdown]
### Output
Returns a .tsv file that reports all possible islands and accompanying information. <br>
Column 1: name of core gene that island is adjacent to <br>
Column 2: length in ORFs of the island <br>
column 3: the indices of the island in the gene neighborhood path of the core gene <br>
Column 4: the names of the genes on the in the gene neighborhood of the core gene. The island can be acessed using the value from column 3. <br>
Column 5: the coverage of genes in the gene neighborhood with the same index correspoinding to column 4.


This file was used as the starting point for visual inspection in Cytoscape.

# %% [code]
for things in primera:
    print("{} \t {} \t {} \t {} \t {}".format(things, derepped2[things], actual_pattern[things + " "], file_paths[things], file_lines[things]), file=open('all_inserts_explore.tab','a'))
```
