## Supplementary material for "Using the pan-genomic framework for the discovery of genomic islands in the haloarchaeon *Halorubrum ezzemoulense*": All of the supplemental material: supplemental_figures_new.pdf

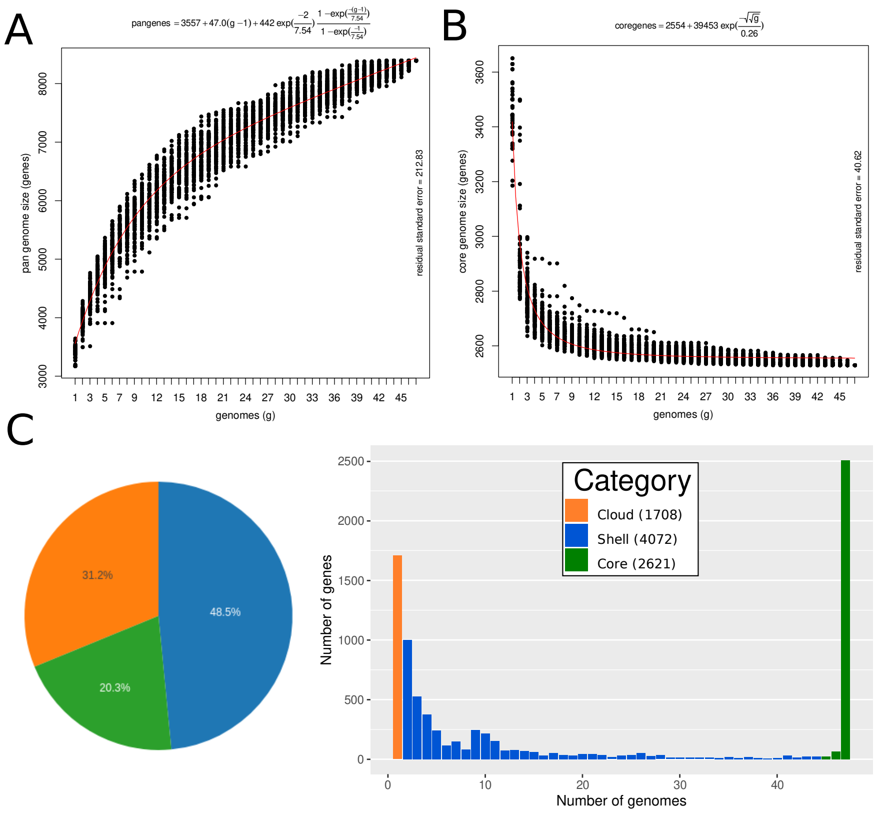


Figure S1**. A description of the *Halorubrum ezzemoulense* pan-genome calculated by Panaroo.** A) Pan-gene accumulation curve as a function of increasing genomes, where there were 100 random permutations per genome increase. The line of best fit is shown in red and fitted to the equation 4 described in Tettelin et al., 2005. This pan-genome is considered open, with a Heap's Law alpha exponent of 0.754 (values >1 is considered closed). B) Core genome decay as a function of the number of genomes sampled. The line of best fit (in red) was calculated using the methods described by Willenbrock et al, 2007. C) Pie chart showing the composition of the core, shell, and cloud genomes, with an accompanying histogram summarizing the distribution of gene family sizes in the dataset.

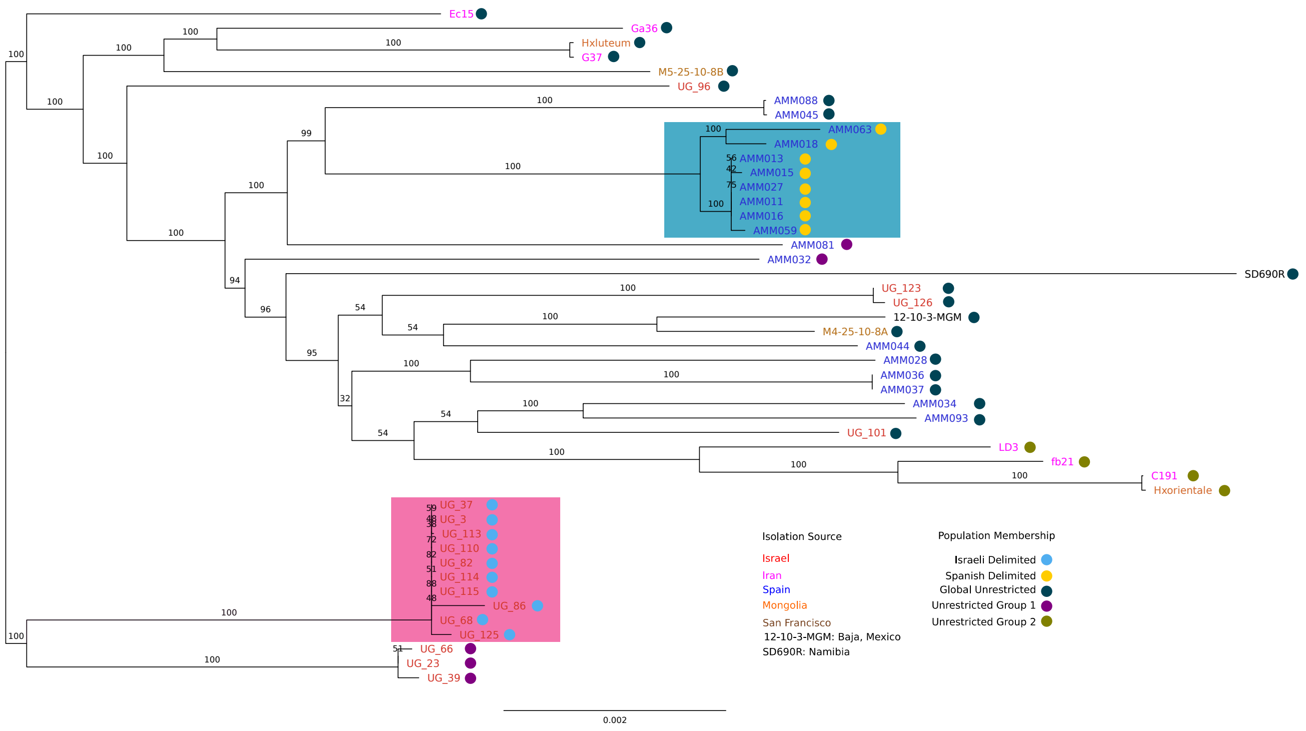


Figure S2. **2,621 core genes phylogeny of *Hez* isolates.** This phylogeny was inferred using a nucleotide concatenate from all of the core genes, using the GTR+F+G4 model in IQTree. The coloring of tip labels indicate isolation location, and the colored circles to the right of the tips indicate population membership when SNPs from neutral core genes were hierarchically clustered into populations. There were 272 core genes with a Tajima's D <|0.2|, indicating they are neutral or nearly neutral. Branch labels indicate ultra-fast bootstrap support (with 1,000 total samples). Most populations group together, with the exception of Unrestricted Group 1, which contains isolates from Spain and Israel.

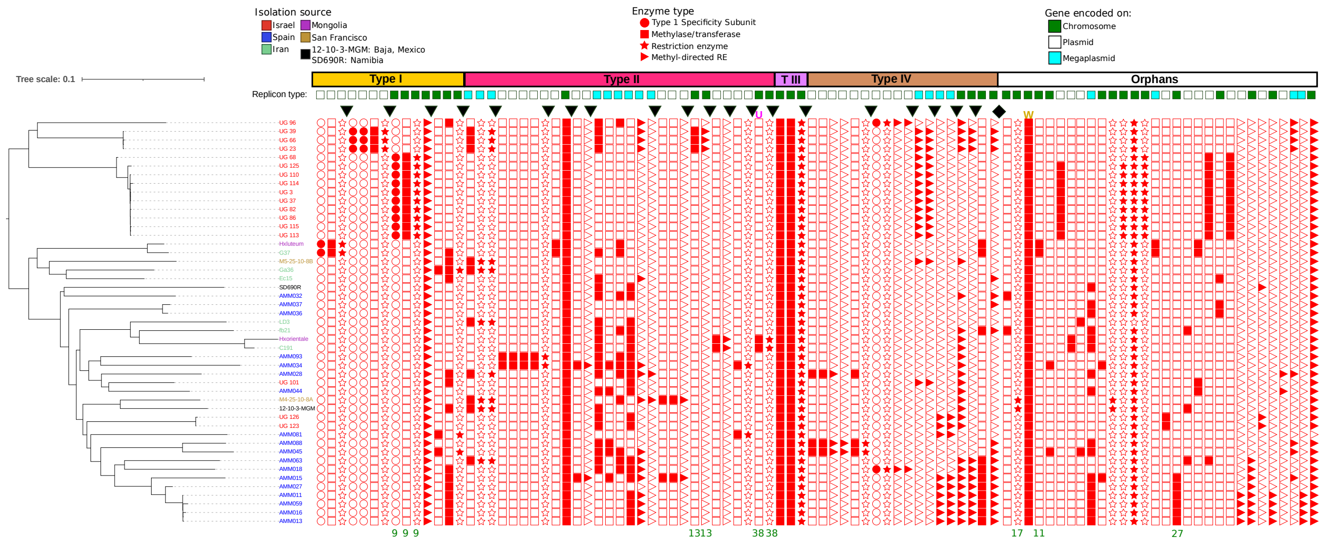


Figure S3. **RMS distribution mapped onto phylogeny**. Restriction modifications systems are sorted first by gene neighborhood membership (±5 genes) then predicted RMS type. Black triangles indicate these gene neighborhoods (i.e., for the first 3 columns those 3 genes are found within the same gene neighborhood, and the same is true of the next 4 genes). Those columns that have a green numbers under them indicate the numbered island that the gene was found on.

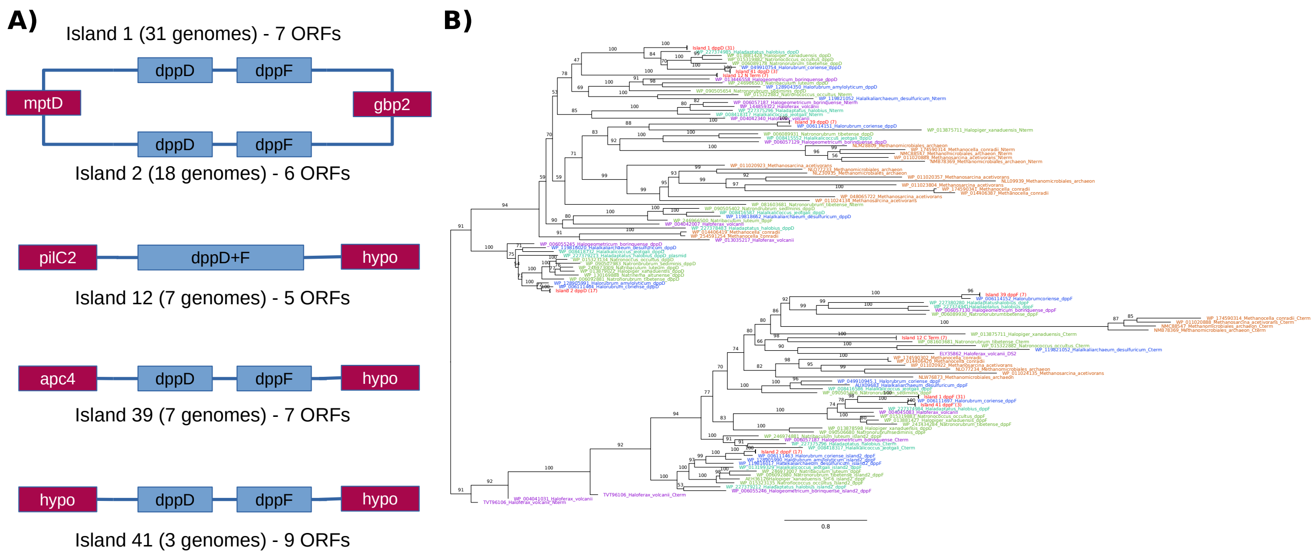


Figure S4. **Variations of the *dppDF* transporters contained on the islands of *Hez.*** A) The various organizations of *dppDF* homologs, and their corresponding invaded core genes (colored in maroon). The other ORFs not related to *dppDF* on the islands have been omitted. Island 12 contains the gene fusion of *dppDF*, whilst the other islands have *dppDF* as separate genes. B) Phylogeny of *dppDF* genes using amino acid sequences of the 9 homologs displayed in A, aligned using mafft-linsi (on default parameters). Island 12 sequences were split in half, with the N-terminal portion corresponding to *dppD* and the C-terminal portion corresponding to *dppF,* and are named as such in the tip labels. The LG+F+R8 model was used to infer this tree, and was determined as the best model using corrected AIC and BIC. Homologous *dppDF* sequences outside of *Halorubrum ezeemoulense* were sampled for each island variant. We broadly sampled other orders of *Haloarchaea* and distantly related Methanogens, including Halorubraceae (blue; which contains *Halorubrum*), Haloferacaceae (purple), Natrialbaceae (light green), Halobacteriaceae (cyan),and methanogens (orange). *Hez* (red) tips were collapsed since there was little sequence divergence between each set. The tree was rooted using the duplication between *dppD* (top half) and *dppF* (bottom half). Branch labels indicate ultra-fast bootstrap support (1000 times).

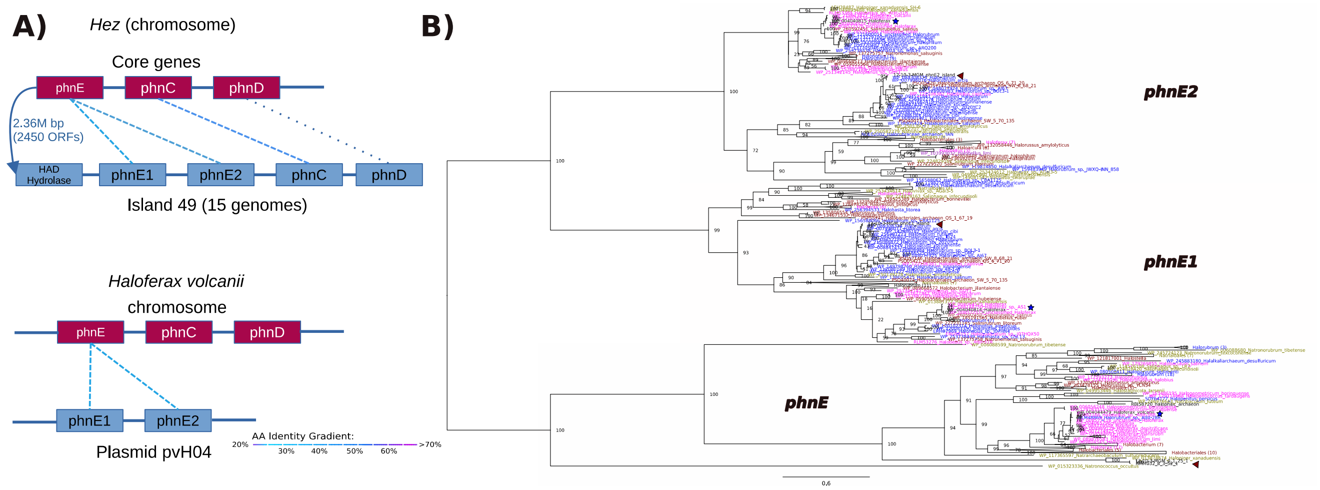


Figure S5. **Comparison of *phnCDE* genes in *Halorubrum ezzemoulense* and *Haloferax volcanii.*** A) Gene schematic of core-*phnCDE* as it is represented in the *Hez* pan-genome graph, and it's homology with the *phnCDE* genes on Island 49. Dashed lines indicate a pair of homologs, colored by amino acid identity. There is an unbalanced gene duplication of the core-*phnE* gene onto Island 49 which leads to two *phnE* on the island. Core-*phnCDE* displays contiguity with the genes encoded on Island 49, in the *Hez* fb21 chromosomal sequence and the pan-genome graph. For contrast, *phnCDE* homologs from *Haloferax volcanii* were drawn. Knockout experiments using the chromosomal *phnCDE* genes determined their association to eDNA uptake (Pearson, 2022). The unbalanced gene duplication of *phnE* also exists in *H. volcanii* (on plasmid pvH04), although no other *phn* genes were found in their neighborhood. The accession numbers for the *H. volcanii* genes are: HVO_A0461, HVO_A0460 (plasmid genes); HVO_0445, HVO_0446, and HVO_0447 (chromosomal genes). B) Phylogeny of phnE homologs with sampling of sequences from Haloarchaea. Tip labels are colored by order and family associations: blue (Haloruburbaceae; closest relatives of *Halorubrum*), olive (Natrialbales), magenta (Haloferacales, excluding Halorubrum) and maroon (Halobacteriales). The brown triangles point to *Hez phnE* sequences, while blue stars point to *Haloferax volcanii* sequences. The LG+F+R7 model was used to infer this tree, and ultra-fast bootstrap values are displayed at the bifurcations (1000 samples).
