## Supplementary material for "Using the pan-genomic framework for the discovery of genomic islands in the haloarchaeon *Halorubrum ezzemoulense*": All of the supplemental material: supplemental_text_mbio_format.docx

Restriction modification systems

RMS genes are typed into 4 different categories (Type I, II and III), and Type IV (1, 2)). Type I-III systems include: a component with restriction enzymatic activity which makes a double stranded cut at a site which is recognized by the enzyme, and a methylase component which prevents restriction activity by attaching a methyl group in the recognition site. Type I systems have a separate specificity subunit (Fig S3). Type IV RMS systems, however, only induce double stranded breaks at target sites which are methylated. These systems are defense systems against foreign DNA (3–5). RMS methylases enable the encoder to mark its own genome as “self” with a methyl group to overcome the cost of endonuclease autoimmunity. Therefore, DNA without a methyl group at the recognition site of the recognition site will be cleaved by the RMS restriction enzyme. However, this defense utility is often limited, as acquisition of an RMS system only decreases the frequency of infection but does not prevent it (6, 7).

While RMS systems are beneficial with respect to defense, diversity generation, and DNA repair (3, 7); their sporadic distributions throughout *Hez* isolates (Fig S3) suggest that some RMS also can be considered selfish genetic elements with the goal of persistence and self-replication (8). Type I-III systems are inherently toxin-antitoxin cassettes which force their way into a host genome through addiction (9). The restriction enzyme (toxin) will cleave the host genome, unless the methylase (antitoxin) prevents it (10). The antitoxin gene product expires before the toxin; which is the basis of the "addiction” the RMS imposes on the host. Thus, the restriction enzyme must be inactivated first for the addiction cassette to be lost. Previous surveys of RMS systems in Haloarchaea revealed most systems are patchily distributed throughout the class (11). In addition, the sporadic distribution of some orphan methyltransferases indicates the loss of the cognate restriction enzyme. Similarly, RMS systems in the *Hez* pan-genome are often found on GIs (Fig S3, Supplemental Spreadsheet S5) or on secondary replicons (Fig S3). Of the 94 RMS related genes found in the *Hez* pan-genome, 88 genes have a limited distribution amongst the 47 isolates, and 30 RMS genes do not have another RMS genes within the 5 gene neighborhood (Fig S3). While the target/recognition site of these discovered systems is impossible to determine based on homology alone, their overall distribution elucidates evolutionary dynamics underlying these systems. While the survival and persistence of these RMS genes contributes to the defense capabilities amongst isolates, their distribution reflects the behavior of selfish genetic elements, that are often lost on the chromosome, but persist within the flexible gene content of the species.

Given the addiction mechanism and the fact that RMS systems are frequently lost or partially deleted and inactivated, these systems may not confer an immediate strong selective advantage to the individual(s) which acquired them so that they become fixed in a population due to selection. However, having a diverse RMS repertoire available in the pan-genome may provide a benefit to the group (i.e., the *Hez* species or population). Varied usage of RMS systems amongst strains of the same population limits rapid virus infection to a subsection of the population which does not have the RMS required to target he infecting virus. This allows for the survival and diversification of the group members which have not been consumed by infection. Additionally, Haloarchaea have a documented history of targeting other Haloarchaea with acquired CRISPRs driving speciation and preventing homogenizing recombination (12). The differential utilization of RMS systems throughout strains of the same species may have a similar effect, erecting barriers to recombination if the methylase does not protect against the restriction of another isolate with a different system.

Restriction modification systems are often recruited to a host or mobile genetic element as a part of a genetic arms race where each participant continuously innovates mechanisms to target the other(s) (13, 14). The target mobile genetic elements are often selected for the ability to incorporate methylase genes to evade the restriction action of the host (5). 7 of 10 island encoded RMS genes are found exclusively on integrase encoding islands (Supplemental Spreadsheet S6), meaning they were not recombined to those islands from chromosomal loci. In addition to selection for transmissibility, there is selection on the recruitment and diversification of different RMS to keep up with the rapid evolution of its genetic predators.

AMM015 – a strain that harbors both islands of a conflicting pair

In the case of strain AMM015, the presence of both opposing genomic islands of a pair is not likely the result of contamination; CheckM (15) detects no contamination or strain heterogeneity (Supplemental Spreadsheet S10). The amino acid divergences between homologs on each island is so great that it is not detected as strain heterogeneity. We calculated the read coverage of opposing islands (Supplemental Spreadsheet S10) in AMM015, and in all pairs: one island had about 100x coverage and the cognate island had ~10x coverage. These results reflect two possibilities: 1) the island with less coverage is present in only 10% of the chromosome copies in AMM015, or 2) the sequencing reaction captured two very closely related strains that only differ in their competitive genomic islands. The former scenario is perhaps made more plausible given the Haloarchaea’s polyploidy, in which they keep ~20-30 copies of their chromosome in a single cell (16), and the coexisting chromosomes may not equivalent (17). One tenth of these chromosomes may have swapped one island for the opposing island, but the island has not been replaced in the other copies.

Solute transporters in *Hez*

Archaeal organisms encode a wide range of ABC transporters (18) including, but not limited to: carbohydrate uptake transporters (CUT) and di/oligopeptide (dpp) transporters (19).These classifications are somewhat misleading however, as CUT transporters often show homology to dpp transporters (19–21). Likewise, dpp transporters are often found in the gene neighborhood of sugar degrading enzymes (22) and some were found to bind sugars (23). The promiscuity of ABC transporter systems to various substrates can be elucidated by their substrate binding protein, and genes found in its gene neighborhood.

Island 19 encodes the only instance of a complete CUT transporter within the *Hez* pan-genome. This island is found in 3 Iranian, 1 Mongolian, and 3 Spanish strains (Fig 2) coexisting in those regions with isolates that do not possess the island. Other components associated with this CUT transporter not encoded on this island, such as malK and malQ are found sparingly in only 2 genomes disassociated with the other components of the transporter (not in the same gene neighborhood). The gene content of Island 19 seems to genuinely resemble a transporter which traffics saccharides (Supplemental Spreadsheet S3). However, it seems to be a chimera of the 2 described CUT operons found in *Pyrococcus furiosus* which import trehalose/maltose (TM system) and maltodextrin (MD system). In addition to the canonical components of an ABC transporter, Island 19 also encodes a fructokinase and the *trmB* regulator found in the TM system, but also includes an alpha-amylase of the MD system (24). The presence of genes from both systems indicates that this CUT cluster may have the ability to transport either substrate. Island 19 is found across multiple geographic regions and its sequences are separated by less than 0.01 substitutions (nucleotide) per site (Supplemental Spreadsheet S7) and has a calculated Tajima’s D value of –1.5. Given its geographic distribution, this locus is more likely to have been under purifying selection, rather than a having gone through a recent bottleneck. The fact that this island is only present in a small fraction of sampled genomes indicates that either it was introduced to *Hez* fairly recently or has limited niche adaptability. The population genetics statistics and the cross regional distribution of the island indicate the former is more likely.

Island encoded transporters reveal a polyamine scavenging mechanism

Of the other 7 ABC transporter islands, Island 16 and Island 49 cannot be readily assigned to either CUT or dpp type islands. Island 16 encodes 5 genes for the import of putrescine/spermidine or polyamines, namely the potABCD system (25). These 5 genes do not have closely related homologs elsewhere in the pan-genome, meaning they were not recombined from other loci onto this island. Island 16 is found in 19 of the 47 *Hez* genomes (spanning geographies), making this a rare but shared utility amongst the species. Polyamines in Archaea perform the covalent modification of translation factor aIF5A (required for translation (26–28)) and the agmatinaylation of tRNA-Ile (which is required for its proper folding (27) ). Island 16 is inserted in between core genes also relevant to translation, nop5 and cbs10 (Supplemental Spreadsheet S9), which suggests the annotated functions of the island are accurate. Early surveys of halophilic Archaea reported the absence of polyamine biosynthesis genes within the entire group of Haloarchaea (29), but these genes were widespread in other Archaea (especially hyperthermophiles). The loss of this capability was presumed to be the result of the salt-in strategy employed by the Haloarchaea, whereby up to 3M intracellular KCL (30) renders the function of polyamines to be redundant or impossible. Later studies of *Haloferax volcanii* revealed the modification of aIF5a was necessary for cell growth, and a putative agmatinase (Hvo_2299) was found in its genome (28). However, it was still unclear how polyamines were synthesized in *Haloferax* in the absence of polyamine synthesis genes. Similar to *Haloferax*, our pan-genomic survey revealed that this agmatinase (*speB*) is a core gene found in 46/47 isolates. This agmatinase, in addition to the genes found on Island 16 presents a possible polyamine scavenging and aIF5a modification mechanism that eluded previous studies. It is unclear why this island is only found in a small fraction of the *Hez* isolates, given its potential utility. Genomes of *Haloferax* have several components of Island 16, but are notably missing the substrate binding protein and one of the membrane permeases. This island is certainly not the only means for halophilic Archaea to acquire polyamines; a mechanism which involves arginine decarboxylases has been proposed to generate agmatine in *Haloferax volcanii* (28)*.* However, those genomes with Island 16 gain a growth advantage in environments with polyamines.

Variations of Trk-type K+ import encoded on a homeocassette

Islands 42+ 43 are homeocassettes encoding TrkAH transporters (Fig 3h), which unlike the other transporter islands is not coupled to ATP hydrolysis (31). These transporters are involved in K+ uptake, which is essential to the salt-in strategy employed by *Hez* (32)*.* Intracellular molar concentrations of K+ is used to counteract the hypersalinity of the external environment and regulate osmotic stress (33). The Trk system is powered by proton motive force and includes a membrane spanning protein (TrkH or TrkG) and a peripheral membrane protein (TrkA) on the cytoplasmic side. TrkA contains a nucleotide binding domain and has been shown to bind NAD+/NADH (31).

Both *trkAH* genes are core genes found in the *Hez* pan-genome and are immediately upstream of the insertion point of Islands 42 +43 (Fig 3h). Both islands encode semi-redundant homologs of the *trkH* gene, accompanied by different accessory genes on either island. We say semi-redundant as the TrkH homologs differ in the number of predicted trans-membrane helices (ranges from 10-12). Interestingly, Island 43 encodes a variant of the *trkA* gene, the other *trkA* gene is found upstream of the island (Fig 3h). The upstream TrkA is 480 amino acids in length, and the island encoded TrkA is 230 amino acids in length and is homologous to either half of the core gene but displays greater similarity to the N-terminal portion. The gene content of these islands closely resembles the archaeal genes from the mosaic hypersalinity island found in *Salinibacter ruber* (33)*.* The *S. ruber* hypersalinity island contains various homologs of the trkAH genes, as well as the universal stress proteins (usp) and amino acid transporters found on Islands 42+43 (Fig 3h, Supplemental Spreadsheet S3). The *S. ruber* island also includes several *trkA* homologs which appear to be half the length of other *trkA* homologs on the island. Notably missing from Islands 42+43, but present in the *S. ruber* hypersalinity isle are Na-K-Cl transporters and *kefB* genes horizontally acquired from Firmicutes and Cyanobacteria.

It was previously thought that the truncated *trkA* gene was the result of a frameshift mutation and was a pseudogene (22). Johnson et al., 2009 cloned and expressed the truncated *trkA* (N-terminal half) along with another truncated *trkA* (C-terminal half, which had lost its nucleotide binding site) from *Thermotoga maritima* into *E. coli* (34)*.* The authors discovered that these two truncated TrkA proteins, along with a novel membrane spanning protein constituted a new sub-class of K+ importers, dubbed the two TrkA importer. The authors expanded this discovery to the two truncated TrkA genes in *Mycobacterium tuberculosis*. However, in this case TrkA in *M. tuberculosis* retained their nucleotide binding domains. This led to the proposal of another sub-class of Trk importers that may not be transporting potassium.

While both Islands 42+43 have the same insertion point and similar gene content, only Island 43 contains the truncated TrkA, which has retained its nucleotide binding motif. It may be possible that the island encoded TrkA can form a complex with the core TrkA (upstream of the island) to create a novel Trk importer. Island 43 also has one more TrkH homolog than Island 42, in addition to the truncated TrkA. Another possibility is that the truncated TrkA forms a complex with the island encoded TrkH, rather than the TrkH found in the core genome. Both of these possibilities and the existence of Island 42, reflect the diversification of potassium import strategies throughout the *Hez* pan-genome. Regardless of which island an isolate possesses, it will still retain its core *trkAH* genes so the fundamental K+ import mechanism is kept in-tact. This exemplifies the diversity possible between two opposing islands of a homeocassette.

Divergent di/oligopeptide transporters are retained in the *Hez* pan-genome on islands

Islands 1 and 2 are competitive pan-genomic islands that both insert between the same core genes (Fig 3a). The term “competitive” refers to the competition for a place between these core genes within a single isolate of *Hez* (as the islands are mutually exclusive on one replicon). The general function for both islands is to encode an ABC transporter for the scavenging of extracellular oligopeptides, also called the *dppABCDF* operon (35). ORFs on opposing islands which encode oligopeptide transport components are divergent homologues (Fig 3a, dashed lines indicate homology across islands), and share less than 50% amino acid identity (as low as 23%). In some cases, there is no significant (<1E-3) nucleotide similarity between homologs, and the amino acid sequences must be used for comparison (blastn, blastp, and PRSS comparisons of opposing island homologs can be found in Supplemental Spreadsheet S10). In addition to the genetic variation of homologs encoded by island pairs, these competitive islands also introduce gene content variation. Island 1 encodes 7 predicted ORFs, while island 2 only encodes 6; only 5 ORFs from each island are related to the *dpp* transporter. The remaining two ORFs from Island 1, and one ORF from Island 2 do not have any detectable homologs on the opposing island; making the gene ensembles on their respective islands unique (Fig 3a).

The case of Islands 1+2 is made more interesting as Islands 12, 39 and 41 also encode variants of the *dppABCDF* transporter. Islands 39 and 41 include 5 ORFs corresponding to the five components of the transporter, however, Island 12 only encodes 4 ORFs (related to *dppABCDF*). It is important to note once again that the di/oligopeptide (dpp) family of ABC transporters is a broad classification, and those systems annotated as dpp transporters may not import peptides at all (19). The propensity of the extracellular substrate binding protein to interact with other ABC transporter complexes results in homology-based annotations, rather than function-based annotations. The annotations of other genes encoded on these islands may shed light on the actual substrate being imported on these islands, however, annotations may not be available for many genes found in this fashion. Thus, the following mentions of dpp transporters refer to its ABC transporter classification rather than the actual substrate each respective system may import.

Islands 1+2 are in the same locus; however, the others (Islands 12, 39 and 41) are integrated in completely different loci, thus only Islands 1+2 are called a homeocassette. One of the accessory ORFs on Island 1 encodes a proteinase (Fig 3a), which hints that this island does indeed import peptides. However, Island 2 does not have this gene nor any of its homologs. One of the substrate binding proteins (dppB) on Island 2 is annotated as a glutathione importer rather than a dpp importer, which hints that this island also imports peptides. Island 12 has one accessory gene, unrelated to the ABC transporter, for which annotations are not available (Supplemental Spreadsheet S3). However, within its 5 gene neighborhood there are several genes related to amino acid metabolism, such as: leuA, ClpP (or related serine protease), and a threonine dehydrogenase. This indicates Island 12 is likely to import peptide substrates. Island 39 includes two mysterious, accessory proteins whose homologs are annotated to encode for mycofactocin biosynthesis (mftE) and cocaine esterase (cocE) (Supplemental Spreadsheet S3). These proteins have been characterized in Bacteria, but their functions are unknown in Archaea. However, homologs of both proteins have been reported to have peptidase activity (36, 37) and are co-distributed in the same locus as a dpp transporter, supporting the idea that Island 39 imports peptides as well. In addition to the dpp transporter on Island 41, there are four accessory genes; three of which have no available annotations. The remaining gene (group_2905) is the membrane component of an ABC transporter, which is reported to import Fe^3+^-hydroxamate (Supplemental Spreadsheet S3; (38, 39)). Interestingly, this gene seems to be recombined from a core gene (found in 44/47 genomes), which shares high identity (88.3%), onto Island 41, while the three hypothetical proteins do not have any highly similar homologs in the core genome. It is unclear what substrate specificities Island 41 has, and not much information can be gleamed from its gene neighborhood which is full of hypothetical proteins. It is probably too speculative to suggest that this is an iron-siderophore uptake system, since there is only one iron related importer gene on the island.

Interestingly, a gene fusion event has combined the two ATP binding proteins (DppD + F) into one gene on Island 12, where the N-terminal portion of the fusion corresponds to the *dppD* gene and the C-terminal half of the fusion is homologous to the *dppF* gene (Fig S4a). The amino acid identities between the 10 homologs range from 20-40%. A phylogeny consisting of all 9 (DppD+F from all four islands) ATP binding proteins (Fig S4b) sheds some light on the history of this protein family. First, *dppF* and *dppD* are clearly the result of a gene duplication, and the sequences from both paralogues are separated into their own subtrees. The subtree of *dppD* can be used to root the *dppF* tree, and vice versa, as it is an ancient duplication. Second, each allele of either paralogue (i.e., Island 1 *dppD,* Island 39 *dppD* are considered alleles of the same paralogue) may have slightly different evolutionary histories, as the topology from each subtree does not mirror the other subtree. Sequences from Island 2 predate the gene fusion event in Island 12 (Fig S4a), but the gene fusion event is found throughout the Halobacterial orders and even in some methanogens. The alleles of Island 1 and Island 41 seem to share a common ancestor in both subtrees. However, the *dppD* gene fusion (Island 12 N-terminal portion) predates the divergence of Island 1 and Island 41 *dppD*, but in the case of *dppF* Islands 1 and 41 predate the C-terminal portion of Island 12. The discordance between the two subtrees is suggestive of gene flow or intragenic recombination events in addition to the divergence accumulated in the sequences.

In four of the seven strains which have Island 39 (AMM032, Ec15, Hxlutuem, and Ga36), the contig which encompasses the island was classified as a plasmid by geNomad (40). However, this classification was accompanied by suspect confidence scores (~ 80%) and low plasmid marker enrichment (< 1.0). In addition, this region was not seen as a plasmid in the 3 other genomes which have Island 39. These results likely reflect the mobile and flexible nature of this locus, which may exist as a part of a plasmid or as integrated into the chromosome. In either case, the local synteny of the genes surrounding the island is kept intact (as that is the unit of the graph-based homology clustering).

These findings suggest that pan-genomic islands serve as reservoirs of genetic variation, and divergent gene cassettes are retained within the pan-genome over evolutionary time scales. The ability of ABC substrate binding proteins to interact with other ABC transporters of a different substrate specificity has been described (41). It is not certain what specific peptide substrate each of the island encoded transporters import, given their low sequence similarity they likely have diverged to accommodate different peptide substrates. This variation on substrate binding proteins allows *Hez* members to adapt to different peptide availabilities in various environments, and the retention of semi-redundant transporters on pan-genomic islands confers niche adaptability through gene transfer withing the species.

The island encoded phosphate transporter is a variation of a homologous core gene neighborhood

Island 49 is an ensemble of 5 genes that can be found in 15 *Hez* isolates that are sporadically found in every geographic region. Surprisingly, every single gene was confidently annotated; together these genes form an ABC transporter for the import of phosphorylated substrates, namely the *phnCDE* operon. Pearson (2022) alludes to the niche adaptability of the *phnCDE* system in *Haloferax volcanii* (Hvo); knockouts of *phnCDE* are unable to utilize nucleotides as a phosphate source for growth (eDNA uptake system; Pearson 2022). This island includes all of the necessary components for the ABC transporter; the permease, ATP coupling protein, and extracellular substrate binding protein are all confidently identified. Additionally, the presence of an HAD superfamily hydrolase on this island, and its reported phosphatase activity (42), affirms the association of this island with phosphate/phosphonate import. In addition to the three components of *phnCDE*, Island 49 encodes a duplicate of the *phnE* gene (permease). Another variant of this operon exists in the *Hez* core genome (i.e., every isolate has it*,* about 2.36 M base pairs (or 2450 ORFs) downstream of the island. The core version of *phnCDE* (core-*phnCDE*) only encodes the three canonical components and does not include the duplicated permease on Island 49 (Fig S5a). Core-*phnCDE* has greater similarity to the chromosomal *phnCDE* genes in *Haloferax volcanii*, for which the knockout experiments were performed (43). *Haloferax volcanii* also has a version of Island 49 on its plasmid pHV4, but this plasmid only contains the duplicated permeases, *phnE1* and *phnE2*, and does not have *phnCD* (Fig S5a). It is unclear what association or interactions the plasmid *Hvo phnE*1+2 genes have, but they do not have corresponding ABC transporter components in their gene neighborhood. In the case of *Hez* and Island 49, the *phnE*1+2 gene neighborhood contains homologous phn transporter components, is not considered a plasmid by geNomad, and displays contiguity with rest of the chromosome.
